# CRISPAltRations: a validated cloud-based approach for interrogation of double-strand break repair mediated by CRISPR genome editing

**DOI:** 10.1101/2020.11.13.382283

**Authors:** Gavin Kurgan, Rolf Turk, Heng Li, Nathan Roberts, Garrett R. Rettig, Ashley M. Jacobi, Lauren Tso, Massimo Mertens, Roel Noten, Kurt Florus, Mark A. Behlke, Yu Wang, Matthew S. McNeill

## Abstract

CRISPR systems enable targeted genome editing in a wide variety of organisms by introducing single- or double-strand DNA breaks, which are repaired using endogenous molecular pathways. Characterization of on- and off-target editing events from CRISPR proteins can be evaluated using targeted genome resequencing. We characterized DNA repair footprints that result from non-homologous end joining (NHEJ) after double stranded breaks (DSBs) were introduced by Cas9 or Cas12a for >500 paired treatment/control experiments. We found that building our understanding into a novel analysis tool (CRISPAltRations) improved results’ quality. We validated our software using simulated rhAmpSeq™ amplicon sequencing data (11 gRNAs and 603 on- and off-target locations) and demonstrate that CRISPAltRations outperforms other publicly available software tools in accurately annotating CRISPR-associated indels and homology directed repair (HDR) events. We enable non-bioinformaticians to use CRISPAltRations by developing a web-accessible, cloud-hosted deployment, which allows rapid batch processing of samples in a graphical user-interface (GUI) and complies with HIPAA security standards. By ensuring that our software is thoroughly tested, version controlled, and supported with a UI we enable resequencing analysis of CRISPR genome editing experiments to researchers no matter their skill in bioinformatics.

## Introduction

The use of programmable, targeted endonucleases has revolutionized the field of therapeutic genetic engineering^1^. CRISPR enzymes form a ribonucleoprotein (RNP) when hybridized with either a 2-part (crRNA + tracrRNA) or a single guide RNA (sgRNA), enabling flexible targeting to genomic loci. With either approach, a short, ~20 nucleotide spacer sequence, which is part of the guide RNA (gRNA), targets DNA with complementarity to the gRNA sequence and introduces a double-strand break (DSB), which can be repaired by non-homologous end-joining (NHEJ) or homology directed repair (HDR)^2^. The NHEJ pathway ligates broken DNA ends and may modify broken ends to find a biochemically favorable ligation product, generating insertions, deletions, and substitutions^3^. The accurate detection and quantification of these editing events at both on- and off-target locations is paramount to ensuring safety for therapeutic applications of CRISPR.

Producing safety information for genome editing therapeutics first involves nomination and interrogation of a set of putative affected off-target genomic loci utilizing *in-vivo*^4,5^, *in-vitro*^6,7^, and/or *in-silico*^8^ methods. After off-target nomination has been performed, alterations in gRNA structure, delivery mechanism, and endonuclease properties can decrease off-target editing effects^9^. Importantly, the use of high-activity and -specificity nucleases^10–13^ in combination with delivery mechanisms that limit nuclease exposure time (e.g. RNP delivery) can reduce off-target editing down to levels that are below the standard Illumina Next-Generation Sequencing (NGS) noise rates^13^. During therapeutic optimization, simultaneous quantification of editing at on- and off-target loci can then be used to expediently determine when sufficient efficacy and specificity has been achieved.

A number of methods have been developed to quantify the population of alleles after editing, including heteroduplex cleavage assays^14–16^, capillary electrophoresis^17^, sanger deconvolution (TIDE/ICE)^18,19^ and next generation sequencing (NGS)^20–23^. Limitations have been described for non-NGS based detection methods, including: limited effective editing range^24^, low sensitivity^25,26^, indel size and type limitations^14,18^, low allelic frequency resolution^26^, and reliance on high quality sanger traces^19,26^. Thus, NGS has become the gold standard for high-throughput accurate genome editing detection^27^, and it is the only method capable of simultaneously quantifying editing at both on- and off-target locations in highly multiplexed samples.

Specialized software tools have been developed to characterize and quantify allelic diversity after a CRISPR experiment from NGS data, but these tools have not yet been comprehensively validated using a genomic scale ground truth^20–22^. These tools generally align NGS reads to a reference sequence by scoring matches, mismatches and missing (gap) aligned nucleotides, selecting the highest scoring of the possible alignments, and annotating allelic variants within a certain distance from the predicted enzyme cut site^20–23^. These tools are challenged by the occurrence of repetitive components in the reference or edited sequences, requiring the algorithm to arbitrarily choose between multiple highly and equally scored alignment options (i.e. secondary alignments), which affect the accuracy of the results^21^. Recently developed tools partially overcome this challenge by prioritizing selection of indel events at the predicted cut site^21,22^, but this approach has not yet been comprehensively validated by examining alleles resulting from Cas9 (blunt cut 3bp from 3’ gRNA end) or Cas12a (two variable nick positions, staggered 4-5bp from 3’ gRNA end) DSB repair events^28–31^.

In this work, we develop a software tool, CRISPAltRations, for the analysis of NGS data generated from amplicon resequencing of CRISPR edited DNA. We characterized the editing profiles of 516 unique on-target guides for two CRISPR-Cas systems: Cas9 and Cas12a. We demonstrate a novel CRISPR-Cas enzyme-specific aligner and optimized application parameters to characterize indel profiles, which together improve the results’ quality. We validate this software tool by benchmarking our software tool against other popular NGS analysis software tools using synthetic NGS data generated to represent 11 gRNAs with a total of 603 GUIDE-Seq^4^ nominated on- and off-target pairs that span a wide variety of genomic sequence features with experimentally modeled indels. Finally, we develop a web-accessible graphical user-interface (GUI) to run CRISPAltRations with cloud resources to empower scientists to securely analyze data and visualize results

## Results

### Iterative characterization and refinement of Cas9/Cas12a editing profiles

#### Software tool iteration 1

To begin, we created a pipeline with no preferential indel realignment, prior to characterization of the positional prevalence and type of edits (i.e., population alleles resulting from DSB repair) induced by IDT Alt-R S.p. Cas9 V3 (Cas9) or IDT Alt-R A.s. Cas12a Ultra V3 (Cas12a) in Jurkat cells (Figure S1, Figure S2). For Cas9 (n=273; average read depth=17,518), indel mutations generally intersected the canonical cut site (median 66% of insertions and 80% of deletions). For Cas12a (n=243; average read depth=7,416), insertions were mostly bounded (median frequency >2%) within a −9 to +2bp window from the PAM-distal nick site (median 3-9% per position) (Figure S2). A median of 84% of deletions overlapped with either the PAM-proximal or distal nick site for Cas12a (Figure S2).

#### Software tool iteration 2

Upon observing that reads containing indels often had equally scored secondary alignments, we performed a round of iterative optimization using our novel position-specific Needleman-Wunsch (psnw) alignment algorithm. We use psnw to re-align the NGS reads described above to the reference sequence using a modified position-specific gap-open/extension vector (scoring vector), which positively scores alignments at or overlapping the cut site or PAM-distal nick site (for Cas12a), similar to previous work^21^ (Figure S3). For Cas9, this increased the prevalence of insertions intersecting the cut site (median 95%), but indels remained bounded at non-canonical cut site positions (Figure S3). For example, a median of 1.8% of total insertion events were bounded −1bp 5’ of the canonical Cas9 cut position. For Cas12a, this increased the prevalence of insertions intersecting the PAM-distal nick site from a median of 7% to 24%. Indels were bounded by positions other than cut sites for both Cas9 and Cas12a, and variability of insertion start positions was higher for Cas12a compared to Cas9 (Figure S3, Figure S4). Cas12a indels were bounded between the two nick sites, and as far as −5bp 5’ of the PAM-proximal cut to +4bp 3’ of the PAM-distal cut. Deletion position profiles for the two enzymes mostly remained the same after this iteration (Figure S3, Figure S4).

#### Software tool iteration 3

With these characterized indel profiles, we further optimized the scoring vector to take a new position-based gap open/extension scoring vector that spanned the entire variant detection window (+/- 20 nucleotides around the cut sites) to select secondary alignments with indels closer to the cut/nick site(s) and indels boundaries enriched in experimental data (Figure 2). This increased indels that were bounded at the −1bp cut position of Cas9 to a median of 2.6% of events (Figure 2A). For Cas9, the majority of insertion events remained at the canonical cut site (median 95%) and −1bp position (median 2.6%), with rare events of insertions at the −2bp (median 0.7%) or +1bp (median 0.4%) positions (Figure 2A,B). For Cas12a, a median of 18% of insertion events occurred at the PAM-proximal nick site. We observed a median of 57% of insertion events did not occur at either the PAM-distal or proximal nick site (Figure 2D). This optimization brought indels closer to the cut site(s), even if the indel may not have been introduced at a canonical cut position.

**Figure 2.**
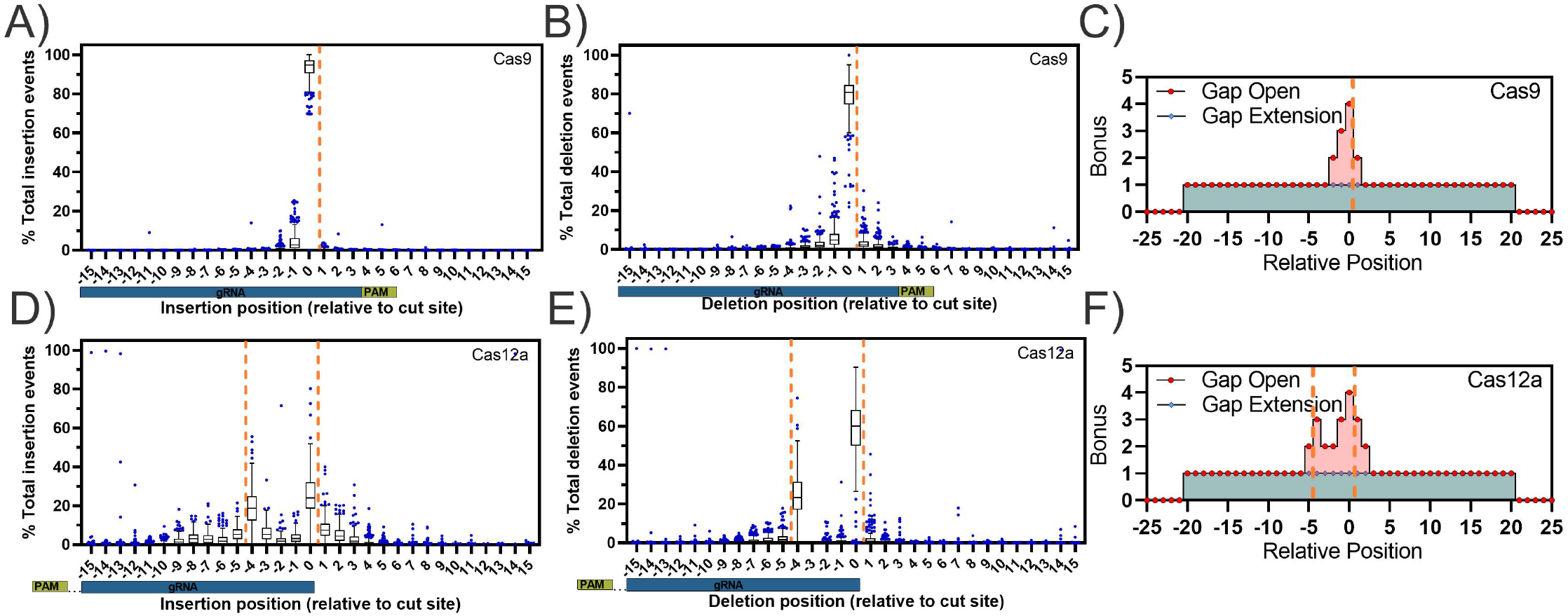
Characterization of Cas9 and Cas12a-specific indel profiles for aligner creation (Software iteration #3). Tukey box and whisker plot of A/D) insertion position and B/E) deletion position relative to the cut/nick site(s) (orange dashed line) derived using C/F) an integrated scoring vector to apply a position-specific bonus to gap open and gap extension events to preferentially select secondary alignments representing the most likely event to occur biologically for Alt-R S.p. Cas9 V3 (n=273 guides) and Alt-R A.s. Cas12a Ultra V3 (n=243 guides) editing events delivered via ribonucleoprotein electroporation into Jurkat cells analyzed using software iteration #3.

### Optimization of the variant detection window limits noise

The variant detection window is a common configurable parameter for CRISPR genome editing quantification software that limits variant calling to a set distance from a predicted DSB, which limits the number of collected false-positive events. To provide a recommended window for quantifying CRISPR editing events in CRISPAltRations, we compared the difference in indel editing events observed in previously used paired treatment and control samples for Cas9/Cas12a in Jurkat cells across a +/- 20bp window from the cut site (or PAM-distal cut site for Cas12a). We determined the optimal window size to be the size at which the median difference of calculated indel editing between treatment and control samples was less than 0.1%. Using this rationale, we find that an optimal window can be defined as +/- 8bp for Cas9 (Figure 3A) and +/- 12bp for Cas12a (Figure S5). However, we found that if the center of the Cas12a window is shifted −3bp from the PAM distal cut site, the optimal variant window can be decreased to +/- 9bp (Figure 3B). Application of this optimal window results in a median decrease in total false-positive indel signal from control samples by 60% as compared to a window size of 20 for both Cas9 and Cas12a while retaining >98% total indel results from treated cells (Figure 3C, Figure 3D). We set these window sizes as the recommended defaults for variant detection in CRISPAltRations.

**Figure 3.**
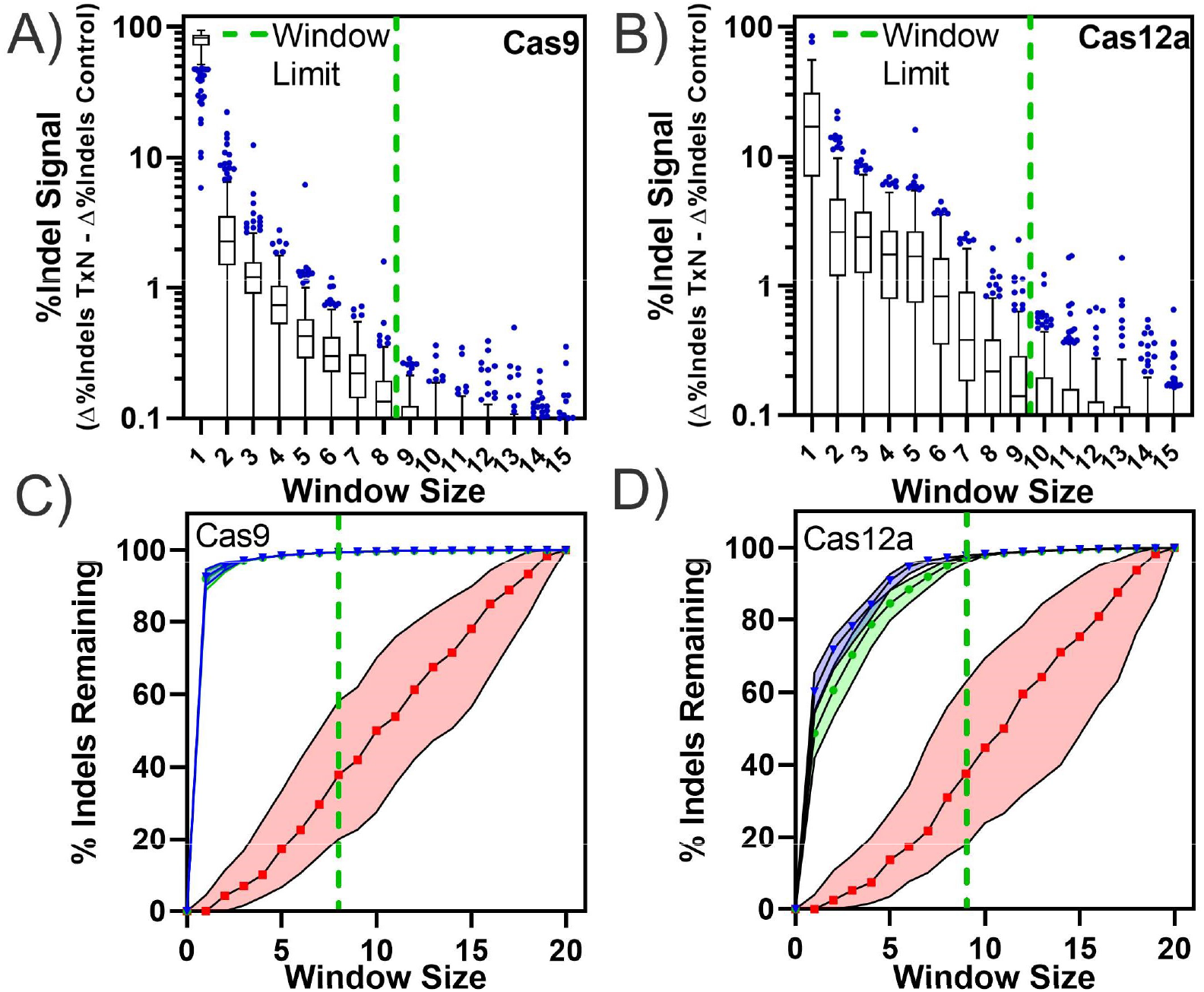
Selection of an optimal variant detection window size. An optimal limit for the variant detection window size (green dashed line) for annotating variants was selected for A) Alt-R S.p. Cas9 V3 (n=273) and B) Alt-R A.s. Cas12a Ultra V3 (n=243; Cas12a window center shifted −3 bp 5’ from PAM-distal nick site) at which median indel signal differences between treatment and control samples was < 0.1%. C/D) The effects of window size on total indels annotated (relative to a window size of 20) was calculated for unedited (red), edited samples with software iteration #2 (green) and edited samples with software iteration #3 (blue).

### Benchmarking of pipeline on- and off-target specificity performance using synthetic datasets

We created a multiplex, synthetic specificity dataset, containing 603 targets, representing performance of 11 gRNAs with indels modeled on observed Cas9 or Cas12a repair events (Figure S6). We created 4,000 synthetic reads per target (50% edited), and we modeled 100 insertion (1-15bp) and 100 deletion (1-25bp) events for a total 120,600 unique indel events (Figure S6). We then validated the performance of CRISPAltRations, and we compared performance with Amplican, CRISPResso, and CRISPresso2.

CRISPAltRations calculated the indel percentage within 0.1% of the expected editing level for 99.5% (600/603) of synthetic Cas9 and Cas12a targets (Figure 4). The three erroneous targets were the result of poor paired-end read merging in regions containing long stretches of homopolymers or repetitive sequence. Observed editing at affected targets deviated from the expected indel percentage by <2% using CRISPAltRations.

**Figure 4.**
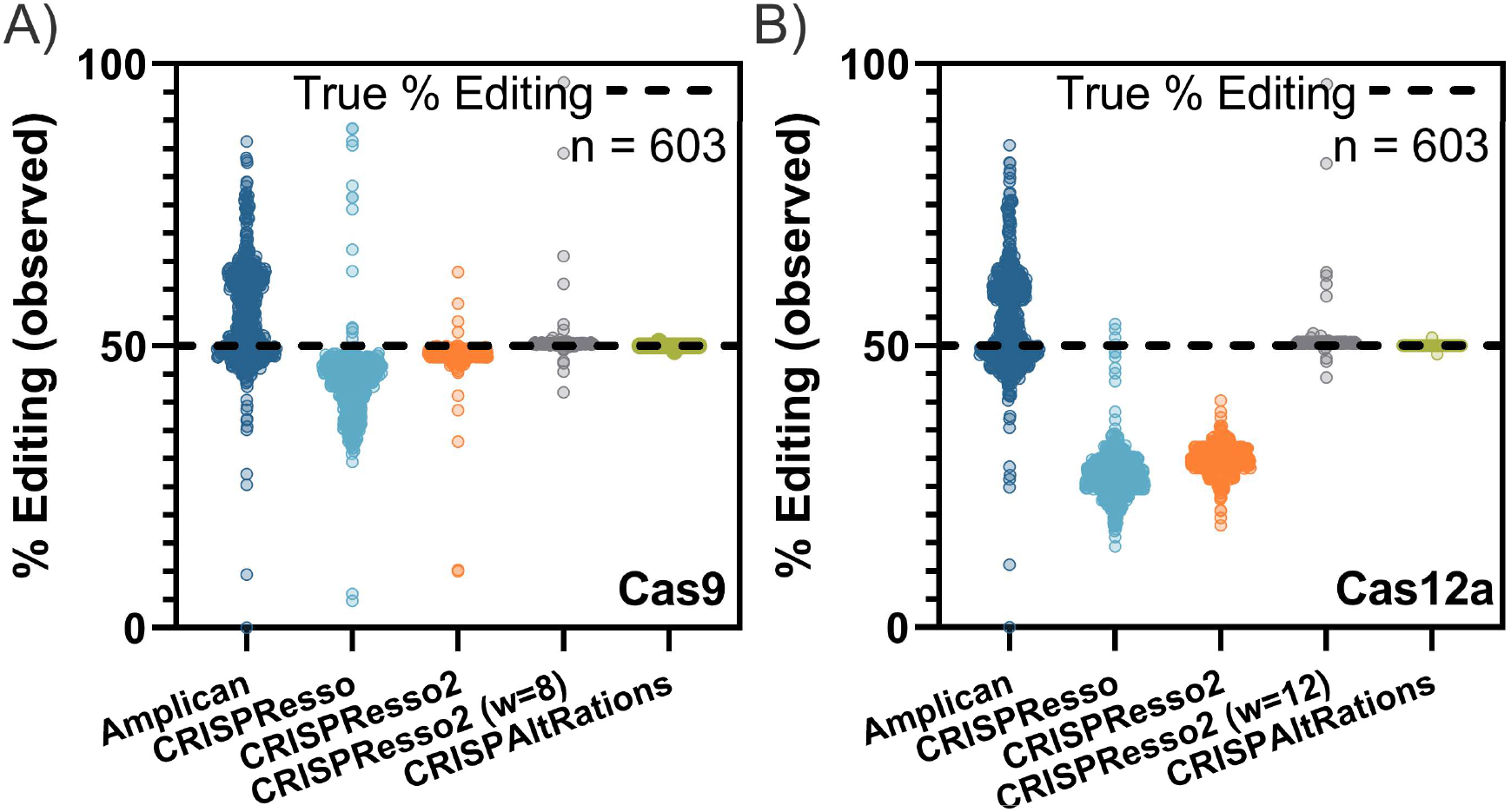
Benchmarking current pipelines supporting multiplex on/off-target analysis. Publicly available tools that easily support multiplex analysis were compared to CRISPAltRations using synthetic data (Figure S6; n=603 sites) generated for A) Cas9 and B) Cas12a for the ability to accurately determine % editing at each site (open circles) with a ground truth of 50% editing (black dashed line). w, *window size*.

We examined the same targets using comparable software tools. The percentage of targets that exceed 2% deviation from the expected Cas9/Cas12a indel percentage for alternative software tools were 72.4%/73.5% (Amplican), 94.5%/99.2% (CRISPResso), 22.4%/100.0% (CRISPResso2), and 1.7%/1.7% (CRISPResso2 with the optimized window parameter derived from Figure 3 and Figure S5) (Figure 4).

### Benchmarking of pipeline on-target HDR accuracy

We created a second synthetic Cas9 on-target dataset (a subset of 91 targets from the previous dataset with equivalent performance between tools) to simulate the performance of the two best performing pipelines, CRISPResso2 and CRISPAltRations, at quantifying HDR rates with a ground truth. This dataset contained each target with a heterogeneous set of events including non-edited events (15%), NHEJ indel events (25%), non-HDR donor integration (15%), imperfect HDR events (15%), and a perfect HDR event (30%). HDR donors were designed to either generate deletions (3, 10, 20, 40bp) or insertions (3, 25, 50, 100bp) within 8bp of the cut site (Methods). The CRISPResso2 software tool was not able to complete data processing on 4 target sites (4.3% total sites) due to an unhandled exception that was not previously present when using the “CRISPRessoPooled” analysis mode on the same sites in the synthetic specificity dataset (Data not shown). These data points were excluded from represented analysis results for CRISPResso2 (Figure 5).

**Figure 5.**
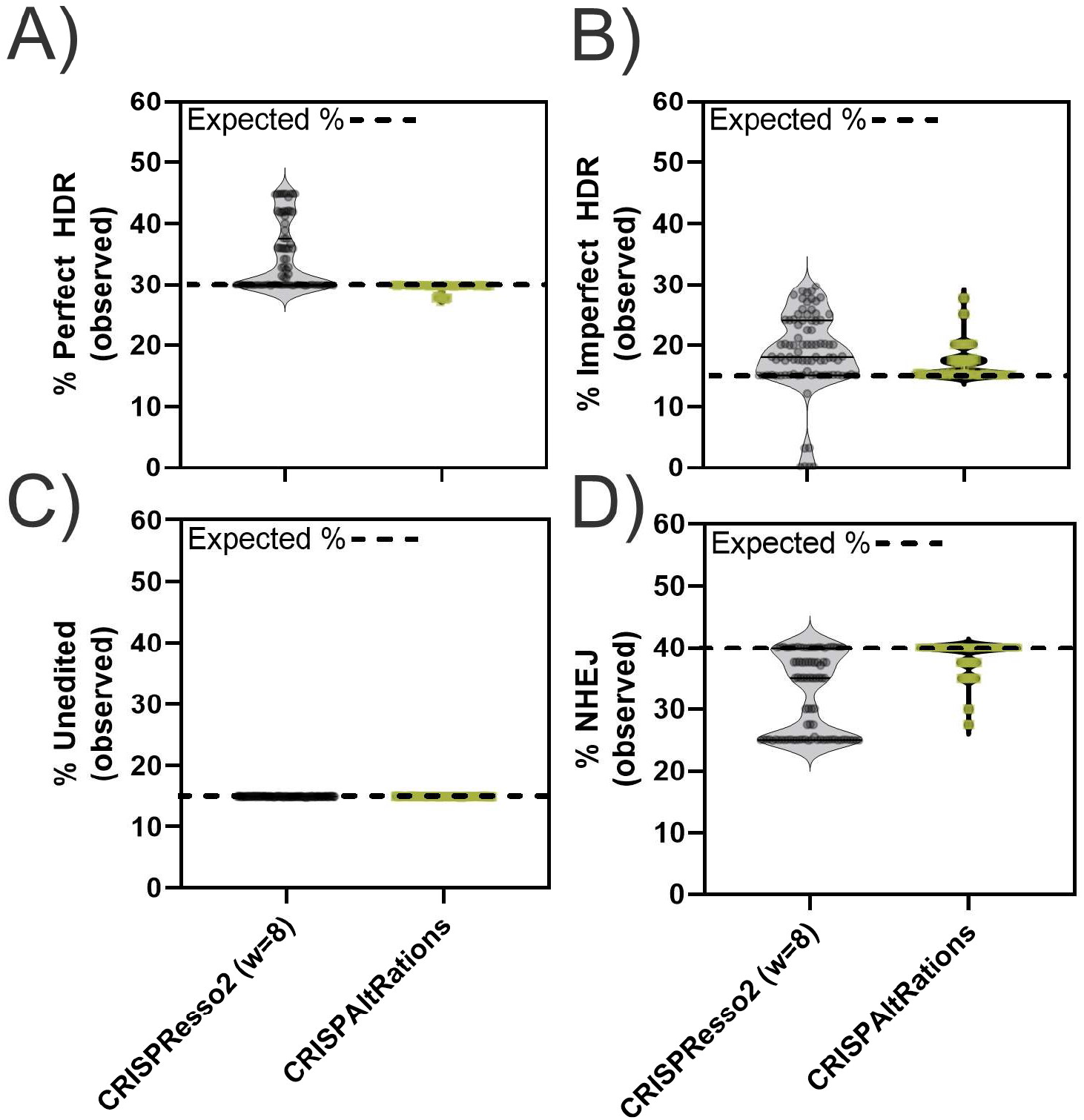
Benchmarking on-target HDR annotation accuracy. CRISPResso2 and CRISPAltRations were compared using a synthetic dataset (n=91 sites) for the ability to accurately determine the percentage of events derived from A) perfect HDR B) imperfect HDR (HDR event with any unintended mutations) C) wildtype and D) NHEJ at all edited sites. w, *window size*.

CRISPAltRations correctly characterized the percent perfect HDR repair at 100% of sites with <2% deviation from truth. CRISPResso2 overestimates the percent perfect HDR repair events by >2% at 43% of sites (Figure 5A). Synthetic HDR-mediated insertions of 50 and 100bp cause the percent perfect HDR of CRISPAltRations to deviate 1-2% below expectation due to the increased probability of SNPs from sequencing errors to occur in these regions (Figure 5B). In contrast, CRISPResso2 does not account for any unexpected SNPs in or near the HDR event in its annotation of percent perfect HDR, which means that any sequencing or polymerase error, naturally occurring mutations, or incomplete HDR events (e.g. 3 out of 4 SNPs successfully incorporated) are not accounted for in its quantification. Both software tools correctly characterize the proportion of CRISPR edited cells at 100% of targets, demonstrating that these differences are not previously identified issues in annotating editing efficiency (Figure 5C).

CRISPAltRations also outperforms CRISPResso2 in its ability to characterize an event as derived from the HDR (imperfect) vs NHEJ pathway at 27 targets (30% of sites) (Figure 5). Overall, CRISPAltRations better characterized HDR editing events in the dataset.

### Using CRISPAltRations to describe mutation profiles of Cas9/Cas12a

We characterized enzyme-dependent (Jurkat/Cas9 vs Jurkat/Cas12a) and cell-line dependent (Jurkat/Cas9 vs HAP1/Cas9) effects on mutation profiles (i.e. indel sizes/types and putative repair pathway) resulting from gene-editing using the improved mutation dissemination present in CRISPAltRations.

Across the 273 targets, Cas9 indel profiles were cell-line dependent. Editing efficiency was >50% in >92% of Cas9 targets for HAP1 and Jurkat cell-lines (Figure S7). The most prevalent mutations in Jurkat cells edited with Cas9 were insertions (median 81%), and a 2bp insertion (median 16%) was the most prominent indel event overall (Figure 6). In contrast, deletions were most prevalent in HAP1 cells (median 75%), and a 1bp insertion (median 18%) was the most prominent indel event overall (Figure 6). Templated insertions (duplication of 1+ nucleotides adjacent to the DSB site) are thought to be a primary mechanism by which insertions are introduced into the genome from repair of DSB events^32^. Insertions in HAP1 cells are predominantly introduced by templated repair events (median 74%). In contrast, insertions in Jurkat cells are introduced by templated repair less frequently (median 8%; Figure 6A). A fraction of insertion events (median 16%) were derived from a non-templated insertion of a repeat of guanine and cytosine nucleotides (GC insertions) of >1bp, an event that did not appear as often in HAP1 cells (median <1%) (Figure 6A). Both cell types derive a fraction of the total deletions from microhomology mediated end-joining (MMEJ) events (deletions with >1bp of exact microhomology; Methods). Deletions mediated by MMEJ were higher in HAP1 (median 43%) compared to Jurkat cells (median 21%; Figure 6A).

**Figure 6.**
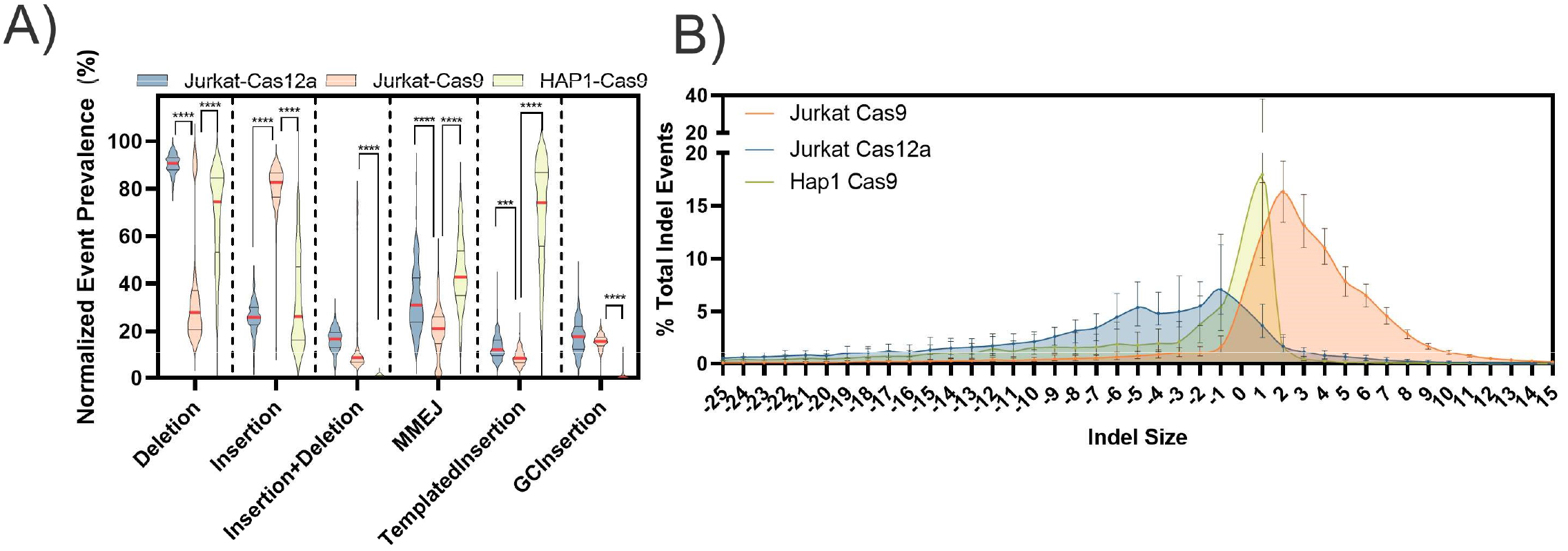
Characterization of cell-line/enzyme specific repair pathways. A) Normalized occurrence of different characterized indel repair events and B) median indel size +/- interquartile range for Alt-R S.p. Cas9 V3 or Alt-R A.s. Cas12a Ultra V3 delivered to Jurkat or HAP1 cells. MMEJ, Microhomology-mediated end-joining.

Comparison of Cas9 targets to the 243 Cas12a targets demonstrated that indel profiles in Jurkat are enzyme dependent (Figure 6, Figure S8). The most prevalent mutation in Jurkat cells edited with Cas12a were deletions (median 90%) and a 1bp deletion was the most prevalent event (median 8%; Figure 6). Insertions mediated by Cas12a editing in Jurkat cells had low frequencies of templated insertions (median 12%). GC insertions were also observed to occur (median 18%) with Cas12a editing (Figure 6A). The normalized abundance of GC insertions was not significantly different (p > 0.05) in Jurkat cells whether Cas9 or Cas12a was used for editing (Figure 6A). DSB repairs mediated by MMEJ were higher with Cas12a (median 31%) compared to Cas9 (median 21%; Figure 6A). Deletion mutations resulting from Cas12a editing were also 6-fold larger than that of Cas9 in Jurkat cells (Figure 6B).

To better understand if mutation profiles could be predicted *a priori*, we compared the spectrum of indels observed to predictions made by *in-silico* repair profile prediction tools, inDelphi^33^ and FORECasT^34^, for all previous targets in Jurkat and HAP1 cells. Both tools perform best when compared to DSB repair events in HAP1 cells with Cas9. In general, FORECasT more accurately predicted the most prevalent mutation, while inDelphi more accurately predicted the spectrum of which indels were observed (Figure S9). For HAP1 cells, FORECasT and inDelphi correctly predict the top mutation event 47% and 41% of the time, respectively (Figure S9B). Both FORECasT and inDelphi predict the outcomes of Jurkat cells edited with Cas9 less accurately, and only predicted the most prevalent mutation type 14% and 10% of the time, respectively (Figure S9B). Both tools predict the repair profiles for Jurkat cells treated with Cas12a (median KL = 0.9) better than Cas9 (median KL = 2.0-2.5; Figure S9A). All predictions made at the canonical cut site of these enzymes are better than those made away from the cut site (−3bp 5’ of cut site) in the same sequence (Figure S9). Predicted frameshift frequencies of both tools correlate with observed results (R^2^ > 0.6), although FORECasT outperforms inDelphi for all cell-line/enzyme combinations (Figure S9).

### Recommendations for experimental read depth requirements and tool limits

We analyzed and subsampled CRISPR NGS data from a series of on- and off-target rhAmpSeq panels (2 panels; 91 and 50 targets) with a wide range of editing frequencies to determine the relationship between read depth and precision. There is an inverse relationship between editing efficiency and number of reads needed to accurately quantify editing (Figure 7). We find that with target coverage >1,000 paired reads per site, >0.5 % indels can be calculated with deviation less than +/- 0.2% indels (Figure 7).

**Figure 7.**
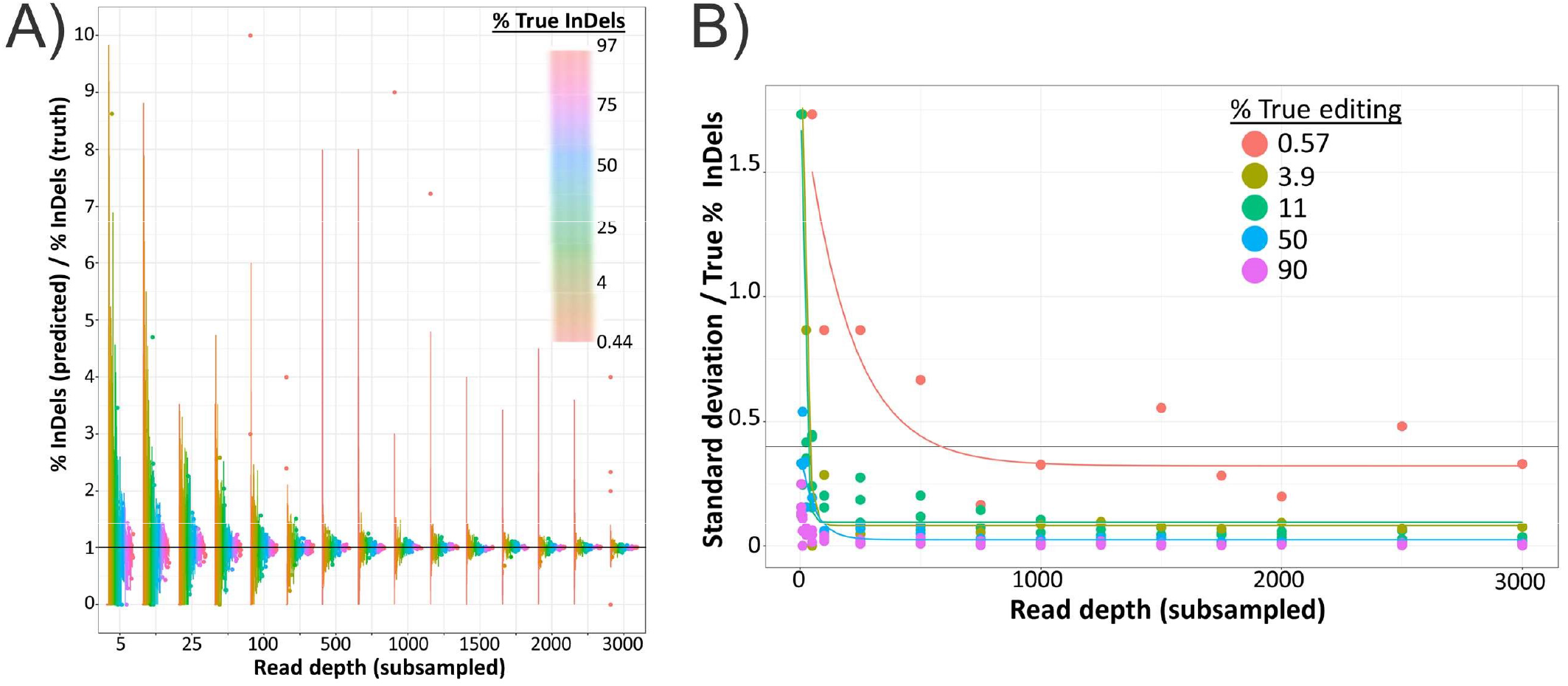
Read depth requirements for variable levels of precision. Subsampling of 284 CRISPR editing experiments with varying editing efficiencies (>0.5% editing) to variable read depths in triplicate with comparison of A) subsampled % indels and B) standard deviation to unsubsampled (i.e., full depth) results.

We evaluated detection limitations using serially diluted DNA standards with rhAmpSeq™ library preparation (IDT, USA) sequenced using Illumina paired-end sequencing. Without any type of background subtraction, the fraction of indels deviated by ~0.2% from the expected standard concentration as indel editing efficiencies approach <1% (Figure S10). After accounting for the indel error rate in a wildtype template using background subtraction, indel editing correlates with expectation (<0.1% deviation) down to 0.1% indel editing (Figure S10).

To better understand the background indel frequencies at diverse genomic loci, we evaluated the indel percentages in unedited control samples at all 273 unique gRNA sites for Cas9 in both HAP1 and Jurkat cell lines. Background indel mutation rates ranged between 0.0-1.0%, depending on genomic locus. Indel mutation frequencies in control samples were found to exceed 0.1% indels ~20% and ~60% of the time for HAP1 and Jurkat cells, respectively. Additionally, for the same set of loci, the limit of blank (LoB) that could be expected in an unedited sample was 2-fold higher in Jurkat cells compared to HAP1 cells (Figure S11). This demonstrates that background indel frequencies can exceed 0.5%, which is above the reported noise rate of Illumina MiSeq instruments^35^ (Figure S11).

### Integration of CRISPAltRations into a cloud platform with a versatile web user-interface

Running computational pipelines can be time-consuming on personal machines and non-intuitive for those unfamiliar with programming interfaces. Thus, we created a web site utilizing cloud-hosted computational resources to run the CRISPAltRations software tool. The web site enables either single or batch file upload of demultiplexed sequencing data files (FASTQ) directly into a cloud-based storage system from a drag-and-drop interface or streamed directly from a sequencer, hard drive, or cloud backup location into the web site. In addition, batch sample analysis is enabled by providing a configuration file (i.e, CSV), and results are summarized in a single report. The web site enables interactive visualization of run metrics including percent editing/frameshift/repair pathway information, percent SNPs for base editing experiments and a heatmap pileup of all allelic frequencies aligned to the reference sequence for visualizing the variant population (Figure 8). In addition to gene editing event summarization, we provide information regarding the performance of the sequencing library and library prep technique used including percentage reads passing QC filters, primer-dimers, uniformity (for multiplex amplification panels) and troubleshooting documents to enable end-users to identify and troubleshoot problematic samples or sequencing runs.

**Figure 8.**
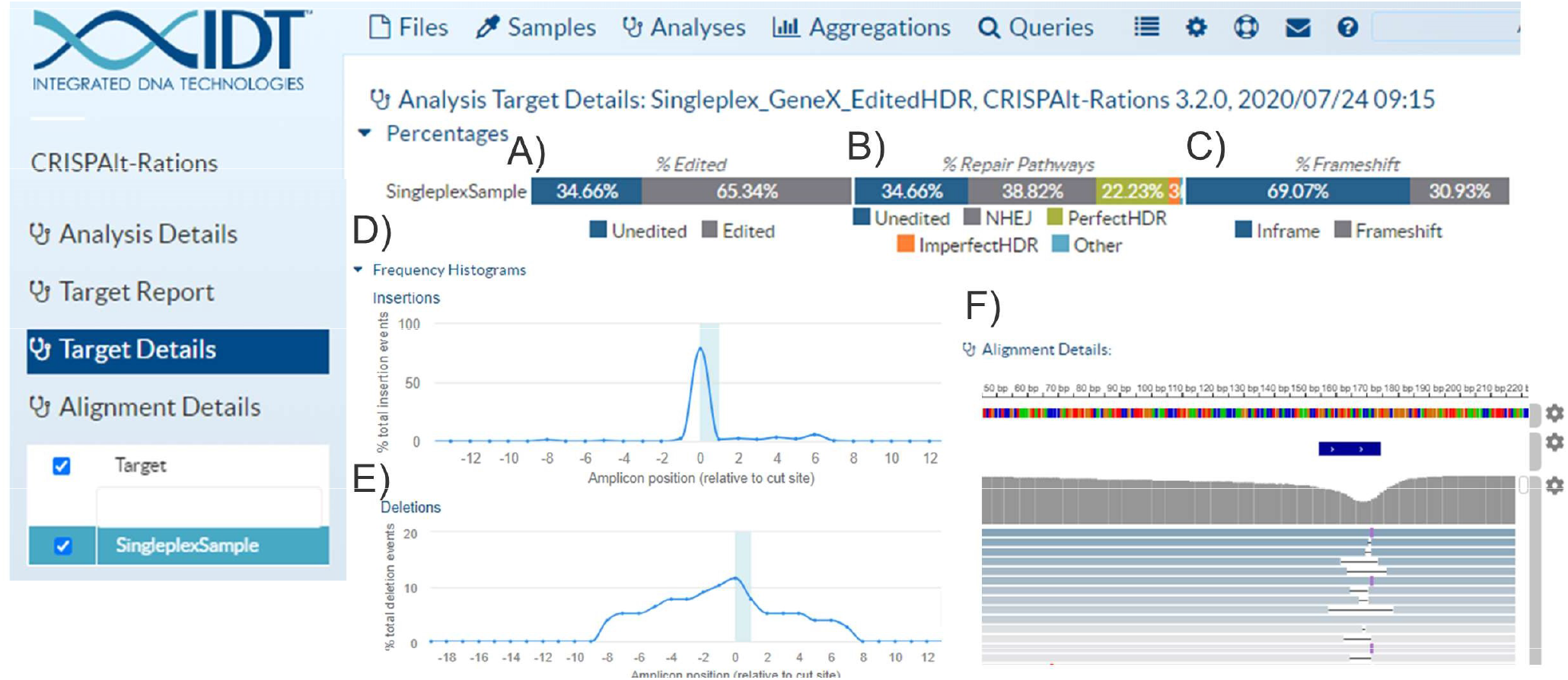
Example of cloud-hosted UI with interactive graphics. As an example, a single on-target HDR experiment is displayed. After completing data processing in the cloud, graphics are automatically created to display high-level metrics like A) Editing frequency B) Repair pathway utilization and C) Frameshift frequency. Additionally, graphics are generated to display positional occurrence of D) insertion E) deletions and F) an IGV visualization of the collapsed variants and their allelic frequencies, and more. Some graphics are artificially condensed to fit in this figure.

We compared runtime performance metrics between CRISPAltRations and publicly available tools processing two synthetic multiplex samples from our on- and off-target benchmarking dataset at various read depth (Figure S12). On common, local hardware, our software runtime is comparable to CRISPResso2 (<40% difference) or outperforms CRISPResso1/Amplican by ~200-750%. Amplican failed time benchmarking on highly multiplexed samples due to a potential unhandled parallelization error (Figure S12). Using the CRISPAltRations web site implementation, runtime is slower (17m to completion) than the local instance on a run with 14 targets (12,000 reads / target), but it remains ~10-fold faster than the CRISPResso2 web site implementation (~4h to completion) (Figure S12). The CRISPResso2 web solution also failed to complete analysis on highly multiplexed (196 targets) or large datasets (>100MB file size) representing an additional limitation (Figure S12). In addition, our web site implementation enables batch runs of thousands of samples simultaneously; while the current CRISPResso2 web site implementation has a maximum of only 4 samples in “batch mode”.

## Discussion

In this work, we develop a software tool, CRISPAltRations, for the analysis of NGS data generated from CRISPR editing experiments. We incorporated knowledge of characterized indel profiles of Cas9/Cas12a into the algorithm, which enhances CRISPR indel detection accuracy. We furthermore show that optimization of the variant detection window reduces false-positive rate, and increases true positive variant calling in Cas9 and Cas12a editing experiments. We benchmark this pipeline against other publicly available, NGS-compatible software solutions using a large synthetic dataset modeled after real Cas9 and Cas12a editing profiles. We demonstrate that our software tool outperforms other available tools. We further demonstrate the utility in CRISPAltRations’ ability to characterize repair profile information, by showing that DSB repair profiles are both enzyme and cell-line specific. Lastly, we provide general experimental recommendations grounded in data for performing CRISPR NGS experiments and access to our tool via a distributed cloud-based web solution with an easy-to-use web site.

Insertions through the NHEJ pathway are primarily introduced at a DSB site. These insertions can be derived from a number of molecular mechanisms including misalignment of microhomologies in cleaved DNA products, staggered overhangs from the cleavage event followed by gap-filling, and/or template-independent polymerase extension^36^. Our quantification of positional insertion prevalence provides unambiguous evidence that insertion events are observed at non-canonical cut site positions, suggesting additional positions that may be subjected to rare endonucleolytic cleavage. It was recently found that Cas9 endonucleolytic cleavage of the non-targeted DNA by the RuvC domain can vary in position relative to the HNH domain cut site to generate a staggered DSB^37^. This in combination with variable degrees of 5’ to 3’ end-processing may explain the positional occurrence of insertions during repair of DSBs introduced by Cas9. For Cas12a we observe a diverse spectrum of positions between +3 bases 3’ of the PAM distal cut site and −5 bases 5’ of the PAM proximal cut site where insertions occur, suggesting a wide range of locations involved in endonucleolytic cleavage and repair. This provides an increased level of resolution on previous work, which has shown that Cas12a cleavage products are diverse and both enzyme and sequence specific^28–30^. This leads us to the conclusion that Cas9 and Cas12a genome editing lead to DSB repair events that cannot be found if only narrow windows (i.e. 1-2 bp) around cut sites are interrogated for variants, a challenge which CRISPAltRations solves with optimized parameter defaults. To the best of our knowledge, this is the first report of the positional prevalence of repair products of Cas9/Cas12a across a wide variety of target sites.

We also demonstrate that indel repair profiles vary with cell- and enzyme-type. Our results support other findings that Cas12a is prone to larger deletions on average when compared to Cas9^38^. Larger deletions have also been shown to be indicative of MMEJ-related repair events^39^. In agreement with this, we find that putative MMEJ events are more predominant in Cas12a deletions compared to Cas9, within the same cell line, suggesting that DSB mechanism contributes to repair pathway preference. Additionally, deletions derived from Cas9 editing in HAP1 cells appear to be more prone to MMEJ than Jurkat cells, suggesting MMEJ prevalence is cell-line dependent due to differences in repair pathway expression/activity. Other mutations such as templated insertions have been reported after Cas9 editing, and they are thought to be the main mechanism by which insertions are introduced during DSB repair^32^. Here we provide evidence that templated insertion prevalence after DSB repair is largely dependent on cell-type, too. The Jurkat cell line has a relatively low frequency of templated insertions, but Jurkat cells had a higher frequency of >1bp insertions containing primarily GC motifs. Future work should address if this type of mutation pattern is widespread in clinically relevant cell-types and identify if it is sourced from a nucleotide bias in a template-independent polymerase. These and other less characterized repair events are poorly predicted in the current generation of *in-silico* indel prediction tools as well, leading to poor performance on Jurkat cells where template-independent mutations are most prevalent. This is likely due to limited repair profile diversity in cell-types used for training these models. In the future, these or new tools could be improved by identifying biomarkers predictive of differential repair outcomes to ensure sufficiently diverse modeling data is generated.

Validation and stability of software has traditionally been an overlooked aspect in bioinformatics program development^40^. Two of the publicly available software tools we evaluated generated uncaught exceptions or run failures at the command-line and web interface on runs that would be reasonably generated for an individual experiment. Additionally, all evaluated software tools were found to inaccurately annotate variants in our benchmarking datasets. Issues resulting in software tool inaccuracies include, but are not limited to, 1) improper target:read assignment, 2) suboptimal read merging strategies, 3) suboptimal alignment strategies, 4) problematic filters/defaults, and 5) general programming errors. Amplican’s performance on this dataset was particularly surprising, and it is primarily caused by the chosen read:target assignment strategy using a string match of the primer binding site based on exact read content. Although we enabled an extra 1bp of ambiguous content (primer_mismatch=1) in an attempt to account for modeled sequencing errors, a fraction of reads were still lost, resulting in inaccurate annotation. Enabling higher amounts of ambiguity in matches leads to increases in memory requirements which can cause the program to crash (Data not shown). CRISPResso1 and CRISPResso2 without an optimized window parameter is mainly affected by the prevalence of CRISPR-associated indel events occurring outside of the default annotation window. Once the annotation window is extended, suboptimal read merging, alignment, and program annotation of variants seem to be primary causes of misannotation.

Previously developed CRISPR NGS software tools have relied on limited synthetic data or focused on experimentally-derived datasets with limited resolution on “truth”, leading to large discrepancies in accuracy of different software solutions. More established applications of variant calling software tools have experienced similar shortcomings, such as for somatic variant calling in cancer genomics^41^, and consortiums/researchers have developed a series of best practices, nomenclature standardizations, and gold-standard datasets for benchmarking software tools^42–44^. With this work we provide a more comprehensive simulated CRISPR NGS benchmarking dataset to identify limitations in analysis software tools and provide evidence that similar best-practices and standards should be established for the genome editing community. In addition, the sensitivity of many of these CRISPR NGS tools have been stated in previous work ranging from 0.01 – 0.1% editing^20,21^. Although we show that detection to ~0.1% indel editing is possible under ideal scenarios, this is a misleading “sensitivity” measurement as it does not account for processes that may introduce variable levels of false-positive editing signals, which may impact reliability in calling variants. This includes variability in methods used for DNA extraction, library preparation, sequencing/technical artifacts, sequence context, and even differences in intracellular milieu, which we demonstrate may be responsible for differences in background editing signal for identical loci in HAP1 and Jurkat cell lines. In other fields, such as cancer genomics, detecting variants even below 5% allelic frequency with high precision/recall is considered challenging^45^. Sophisticated methods incorporating unique molecular identifiers (UMIs), paired treatment/control background subtraction, and more have all been applied within the cancer genomics field to enable high specificity detection of variants at sub-1% allelic frequencies^46,47^. We highlight here for CRISPR NGS analysis that even background editing signal can vary dramatically based on experimental conditions, further emphasizing the need for statistical tests, replicates and other advanced methods for confident detection of low editing levels. Future work will need to incorporate error-correction sequencing strategies (e.g. UMIs) and more sophisticated background subtraction methods to increase accuracy of editing annotation.

As genome editing therapies enter clinical trials, it becomes a necessity that software and sequencing methods are thoroughly vetted to prevent incorrect conclusions or exclusion of variant information. This has become clear with accumulating evidence that dsDNA donor (e.g. plasmids, AAV) integrations^48,49^, translocations^50^, and large indels/rearrangements^51^ all take place from DSB mediated genome editing. We show that for small dsDNA donors, CRISPAltRations more accurately discriminates and quantifies NHEJ, imperfect HDR and perfect HDR than existing pipelines using simulated data. However, detection of many larger events requires advances in the use of long read sequencing and targeted hybridization/capture-based sequencing, enrichment protocols, and analysis tools. By testing, versioning, and deploying CRISPRAltRations within a cloud-hosted user-interface with reproducible code production environments and security certifications, we aim to provide a plug-and-play hardware-independent solution to generate high quality genome editing specificity data.

## Methods

### Ribonucleoprotein complex formation

Cas9 gRNAs were prepared by mixing equimolar amounts of Alt-R™ crRNA and Alt-R tracrRNA (Integrated DNA Technologies, Coralville, IA, USA) in IDT Duplex Buffer (30 mM HEPES, pH 7.5, 100 mM potassium acetate; Integrated DNA Technologies), heating to 95°C and slowly cooling to room temperature or using Alt-R sgRNA (Integrated DNA Technologies) hydrated in IDTE pH 7.5 (10 mM Tris, pH 7.5, 0.1 mM EDTA; Integrated DNA Technologies). Cas12a gRNAs consisted of Alt-R™ Cas12a crRNAs (Integrated DNA Technologies) hydrated in IDTE pH 7.5. RNP complexes were assembled by combining the CRISPR-Cas nuclease (Alt-R S.p. Cas9 Nuclease V3 or Alt-R A.s. Cas12a Ultra V3; Integrated DNA Technologies) and the Alt-R gRNA at a 1.2:1 molar ratio of gRNA:protein and incubating at room temperature for 10 minutes. The target specific sequences of the gRNAs used in this study are listed in Table S1 for Cas9 and Table S2 for Cas12a. The guides chosen were either within the same general genetic context (same amplicon sequencing space; enzyme-dependent) or identical between the two cell lines (cell-line dependent) used in this study.

### Cell culture

HAP1 cells were purchased from Horizon Discovery (Cambridge, UK). Jurkat E6-1 cells were purchased from ATCC® (Manassas, VA, USA). Cells were maintained in RPMI-1640 (Jurkat) or IMDM (HAP1) (ATCC), each supplemented with 10% fetal bovine serum and 1% penicillin-streptomycin (Thermo Fisher Scientific, Carlsbad, CA, USA). Cells were incubated in a 37°C incubator with 5% CO_2_. HAP1 cells were used for transfection at 50-70% confluency. Jurkat cells were used for transfection at 5-8 x 10^5^ cells/mL density. After transfection, cells were allowed to grow for 48-72 hours in total, after which genomic DNA was isolated using QuickExtract™ DNA Extraction Solution (Epicentre, Madison, WI, USA). We chose HAP1 and Jurkat since they are derived from human chronic myelogenous leukemia and T lymphocyte cell lines, which are derived from cell types that are similar to those that have been best studied in the context of predicting Cas9 repair profiles^33,34,52^.

### Delivery of genome editing reagents by nucleofection

Electroporation was performed using the Lonza™ Nucleofector™ 96-well Shuttle™ System (Lonza, Basel, Switzerland). For each nucleofection, cells were washed with 1X PBS and resuspended in 20 μL of solution SF or SE (Lonza). Then, cell suspensions were combined with an RNP complex. For Cas9, the RNP concentration was 4 μM with 4 μM Alt-R Cas9 Electroporation Enhancer. For Cas12a, the RNP concentration was a suboptimal dose of 0.2 μM with 3 μM Alt-R Cas12 Electroporation Enhancer (Integrated DNA Technologies) to provide a more diverse range of editing frequencies. This mixture was transferred into one well of a Nucleocuvette™ Plate (Lonza) and electroporated using manufacturer’s recommended protocols. After nucleofection, 75 μL pre-warmed culture media was added to the cell mixture in the cuvette, mixed by pipetting, and 25 μL was transferred to a 96-well culture plate with 175 μL pre-warmed culture media. Transfection plates were incubated at 37°C and 5% CO2.

### Quantification of editing by next-generation sequencing (NGS)

On-target editing efficiency for Cas9/Cas12a nucleofected cells was measured by NGS. Libraries were prepared using a previously described rhAmpSeq amplification-based method^53^. Briefly, the first round of PCR was performed using target specific primers. A second round of PCR was used to incorporate P5 and P7 Illumina adapters to the ends of the amplicons for universal amplification. Libraries were purified using Agencourt® AMPure® XP system (Beckman Coulter, Brea, CA, USA), and quantified with qPCR before loading onto the Illumina® MiSeq platform (Illumina, San Diego, CA, USA). Paired end, 150 bp reads were sequenced using V2 chemistry. Data were demultiplexed using Picard tools v2.9 (https://github.com/broadinstitute/picard).

### CRISPAltRations algorithm

We developed the CRISPRAltRations software tool in python, and it plus other software tools are together managed by a snakemake or CWL workflow manager (Figure 1) ^54,55^. The software is hosted with a front-end graphical user interface (UI) at (https://idtcrispr.bluebee.com/idtcrispr/#!login). The UI enables the end-user to specify run information, which is used to partition computational resources hosted in the cloud to perform all data processing using the CRISPAltRations software tool. Results can be visualized and downloaded from the UI. Sequencing data stored in the cloud (AWS, BaseSpace, Google) or on local data stores can be automatically synced with the platform using or uploaded through a “drag-and-drop” mechanism within the UI. Data are processed in region specific data centers, duplicated, and protected in a manner that is GDPR, HIPAA, DSPT, PHIPA, PIPEDA, and CSL compliant.

**Figure 1.**
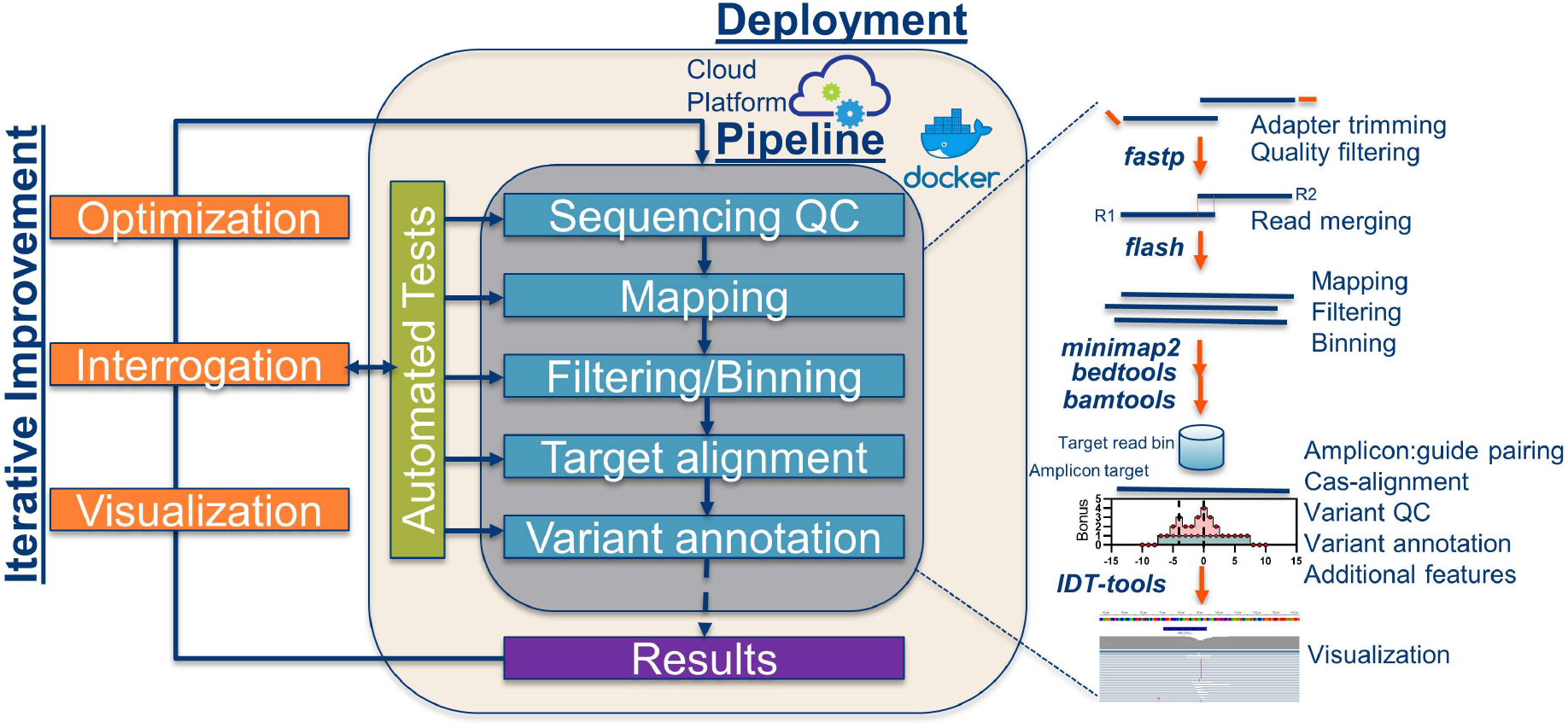
Development framework for CRISPAltRations. CRISPAltRations was architected such that each step of the pipeline (grey box) is containerized and deployed within the cloud to enable highly scalable batch processing (tan box). Briefly, the pipeline goes through a number of processing steps (blue boxes) to transform demultiplexed reads to results that quantify editing events after CRISPR genome editing (purple box) which can be viewed and stored in the cloud or downloaded locally. To improve CRISPAltRations, we used iterative improvement (orange boxes) to iterate through a process in which we manually inspected and interrogated experimental results to build tests (green box) which ensure stability, coverage of different experimental use-cases, and allowed us to optimize the software tool.

The CRISPAltRations software tool workflow starts from demultiplexed FASTQ files as input along with guide and amplicon information in the form of strings or six-column BED-formatted genomic coordinates. The pipeline assumes that the end-user has generated Illumina sequencing data (single or paired-end) in FASTQ format and that the reads completely span the cut site in both directions after merging of R1/R2 pairs. If genomic coordinates are provided in BED file format, amplicon and guide sequences are extracted from the selected genome and paired using bedtools^56^. Next, low quality reads and Illumina sequencing adapters are removed using FASTP^57^ (--adapter_sequence=AGATCGGAAGAGCACACGTCTGAACTCCAGTCA; -- adapter_sequence_r2=AGATCGGAAGAGCGTCGTGTAGGGAAAGAGTGT; -L; -n=10; -q=15; -u=30). If paired-end data were used, read pairs are merged into a single fragment using FLASH^58^ (-O flag used). Putative primer dimers are identified based on a size limit (<60bp reads), annotated by homology to known amplicon sequences, and removed from downstream analysis. The remaining reads are then mapped to all potential amplicon targets using minimap2^59^ (default parameters). The mapped reads are separated into amplicon target-specific BAM files using bamtools^60^ to enable parallel processing of all targets. If an HDR donor was supplied, the theoretically perfect HDR event is recreated by iterating through a Needleman-Wunsch alignment with a high gap-open penalty implemented in biopython^61,62^ (match=2; mismatch=1; gap open=-30; gap extension=0) at all potential amplicons, choosing the optimal query:target assignment, reconstructing the hypothetical sequence based on the alignment and adding the hypothetical sequence to the mappable amplicons reference file. Reads are collapsed based on exact sequence identities and re-mapped to the mappable amplicons reference file using minimap2^59^ (A=2; B=4; O=8; E=5; --secondary=no; --no-end-flt; --max-chain-iter 100000) to bin reads appropriately between events derived from HDR vs NHEJ repair pathways. Mapped reads containing indels are re-aligned using a modified Needleman-Wunsch algorithm we call psnw (https://github.com/lh3/psnw) that attributes an alignment score bonus to placement of gap-open or extension in specific locations in the alignment. Psnw extends the features of Needleman-Wunsch to include an elevated match/mismatch/gap open/gap extension scoring matrix (multiplied by a scalar) and a customizable position specific gap-open/extension vector giving a configurable bonus to alignments that place these features in specific positions. The scoring matrix enables the algorithm to select alignments that have gap open/extensions at desired positions. All reads with a mutation that begins within a set distance from the predicted canonical cut site(s) are annotated and summarized in the results, with a number of other visualizations and reports.

### Variant annotation

Annotation of variants is performed in a step-wise process with custom python code. First, variants are further collapsed based on their annotated nucleotide changes within range of the cut site window. Then, if an HDR donor is supplied, a variant is determined to be derived from the HDR vs NHEJ repair pathway based on the reference amplicon that the read mapped to (wildtype vs theoretical HDR event). Next, a variant is annotated as an imperfect HDR event if any SNP or indel is found within the pre-defined window from the cut site or from the location of the first mutation incorporated from the HDR event to the last, whichever is larger. Next, insertions, deletions, and insertion+deletion frequencies are quantified relative to the reference sequence.

Insertions are then further characterized by inspecting the sequence of the insertion and surrounding genomic context. If the sequence of an insertion is found to be an exact repeat of DNA adjacent to its insertion, it is described as a templated insertion^63^. If the sequence of an insertion is not found to be a templated insertion, and it is found to be composed of >1 nucleotide and contain only guanine/cytosine nucleotides, it is described as a GC insertion. These events are represented as percentages of the total number of insertions to enable easy comparison between targets.

Deletions are further characterized by inspecting surrounding genomic context of the deletion. If a deletion is >1 nucleotide in length and found to contain >1 nucleotide of exact microhomology from the start of the deletion to the 3’ end of the remaining genomic sequence or from the end of a deletion to the 5’ end of the remaining genomic sequence (accounting for secondary alignments), it is annotated as a MMEJ event. MMEJ events are represented as a percentage of the total number of deletions to enable easy comparison between targets. Any events with both insertions+deletions are excluded from this analysis.

Indel mutations that are not multiples of 3bp are annotated as frameshifting events, independent of whether they intersect known coding sequences. For identification of the position of mutations, an insertion position is described as the 5’ reference base position adjacent to the insertion. For deletions, the position is considered to be the position closest to the cut site at which a reference base is missing. Additionally, a deletion was considered to intersect the cut site if the base directly 5’ of the cut/nick site was missing in the variant. Since the cut site(s) of A.s. Cas12a Ultra V3 with a 21bp spacer have not been explicitly defined, we annotated the PAM-proximal and PAM-distal nick sites to be the position between the sites where the most insertion events were observed prior to algorithm optimization.

### Synthetic read generation for on- and off-target editing validation

To create a synthetic benchmarking dataset reminiscent of CRIPSR editing, we used VarSim^64^ for generating the defined variants in a paired-end amplicon sequencing read format with an Illumina MiSeq v3 error profile and ART^65^ to generate unmodified reads with MiSeq v3 error profiles to enable addition of “wildtype” reads with desired error-profiles. We used this to generate a synthetic dataset using sequence space from 11 real rhAmpSeq panels (Table S3) representing GUIDE-seq nominated Cas9 on- and off-target sites (n = 603 on- and off-target sites) with indels modeled based on our real Cas9/Cas12a editing events in Jurkat cells. To do this, median mutation size, position, and frequency of event types across these two datasets were used to create a series of mutation probability vectors that describe the probability of observing different editing events relative to the canonical cut site in a random guide. To create indels, mutation probability vectors were sampled to create 100 unique insertion and deletion events for each guide, each unique event with a read depth of 10 (4,000 reads per target; 50% indels; 2×150 reads). It should be noted that the Cas12a sites are not true experimentally determined Cas12a off-targets or binding sites, but were merely created at the same genomic positions as the Cas9 dataset to recapitulate the challenge to bin reads between on- and off-target sites with similar genomic context.

### Synthetic read generation for on-target HDR quantification validation

To create a synthetic benchmarking dataset representing the ability to perform on-target HDR quantification, we took all of on- and off-targets from the RAG1 Cas9 GUIDE-Seq panel and separated these out as single targets (91 total)^25^. The RAG1 panel was chosen because 1) no target processing problems were found when using CRISPResso2 and 2) the genomic sequence around the targets included homopolymers and other events that represent challenging genomic regions to annotate. We then created dsDNA donors *in-silico* (as sequence strings) with 40bp homology arms using the same synthetic generator previously described. Donors were designed to synthetically introduce a mutation at each of these sites as a deletion (3, 10, 20, 40bp) or insertion (3, 25, 50, 100bp) within 8bp from the expected cut site. We modeled the dataset with simulated dsDNA donors since this introduces an additional potential complication of the actual donor sequence being directly ligated into the cleavage site which is an important event to discriminate between^4,48^. We made all sites have a heterogeneous set of events including non-edited events (15%), 10 unique NHEJ indel events (25%), 5 unique non-HDR donor integration (15%), 5 unique imperfect HDR events (15%), and 1 perfect HDR event (30%). NHEJ indel events were modeled using the mutation probability from Jurkat with Cas9 (see above). Integration of the donor (non-HDR donor integration) was modeled with one perfect integration of the complete dsDNA donor at the cut site and 4 imperfect integrations. Imperfect integration events were modeled with random sizes of truncations of the integration event (not to exceed 40% the full dsDNA donor size) or SNPs within the integrated donor. Imperfect HDR events were similarly modeled with either truncated events (deletion or insertion HDR events) or SNPs (insertion HDR events) within the portion of DNA that was intended to be altered by the HDR donor. Reads were simulated with MiSeq v3 noise profiles (4,000 reads per target, 2×250 reads).

### Determination of required read depth levels

To provide recommendations for target sequencing read depth requirements, we re-analyzed previously published CRISPR NGS data from a series of rhAmpSeq panels designed for on/off target sites of guides targeting the RAG1/RAG2 loci with a wide range of editing frequencies, obtainable at the Sequence Read Archive (SRA) under: PRJNA628100^25^. Reads from these samples were subsampled, without replacement, in triplicate with random seeds to a range between 5-3,000 reads pairs per site and quantified using CRISPAltRations with optimized parameters. Indel frequencies and standard deviation between all three read depth replicates were then compared to the frequency obtained using all reads for the corresponding on- and off-target site to determine deviation from expectation.

### DNA standard titration for evaluating rhAmpSeq accuracy

Synthetic dsDNA templates were generated as gBlocks (Integrated DNA Technologies, USA) using simulated events at an HPRT Cas9 genomic locus (Table S4). Templates were quantified using qPCR before being pooled at equimolar concentrations. These synthetic events consisted of 10 deletions, 10 insertions, and 3 SNPs spiked in to create a known mixture (43.5:43.5:13). Serial dilution was performed with varying levels of wildtype sequence ranging from 0-100% (Table S4) and subjected to the previously stated library preparation procedure followed by NGS.

### Statistical and Data Analysis

Data collected from experiments were analyzed and statistics generated using Graph PadPrism 8. Editing data for Cas9/Cas12a experiments were only used if a sample had >100 merged reads obtained, and the treated sample had >5% editing. Significance was evaluated using a 2-way ANOVA with a *post hoc* Tukey multiple comparisons test (*p < 0.05, **p < 0.01, ***p < 0.001, and ****p < 0.0001) for indel profile differences between Cas9 (Jurkat), Cas9(HAP1) and Cas12a(Jurkat) treatments. Limit of blank (LOB) was calculated using methods previously described^66^.

### Software versions and parameters utilized

For benchmarking analyses the following softwares and versions were used: CRISPResso (1.0.13), CRISPResso2 (2.0.40), Amplican (1.6.2). When using Amplican, the following non-default parameters were used: average_quality=15, min_quality=1, primer_mismatch=1, min_freq=0.000001. For comparison of in-silico repair profile prediction tools the following software versions were used: inDelphi (commit tag: 9ab67ca53ebb91e49aeb4530ec1e999ee9827ca1) and FORECasT (commit tag: 019a2f52ba8437528298523c79c224c205146f00). For both models, the “K562” model was used for comparing performance.

## Availability

The CRISPAltRations pipeline is available via a cloud-hosted web UI at https://idtcrispr.bluebee.com/.

We provide subscription models to cover regular cloud computing usage costs, or provide the interface free-of-charge to customers utilizing rhAmpSeq products for generating their sequencing libraries. Credits to enable a trial of the service can also be obtained by contacting. The psnw aligner is available at https://github.com/lh3/psnw. All Cas-specific gap-open/extension scoring vectors (for psnw) and parameters for publicly available tools are disclosed in Methods for reproducibility.

## Acknowledgements

We would like to thank the Molecular Genetics group at IDT for many helpful discussions. We would also like to thank all of the individuals who participated in testing phases of the CRISPAltRations web platform and their helpful feedback that led to an ultimately better user-interface.

G.K, A.J., M.M., R.T., G.R., N.R., H.L., L.T., Y.W. and M.B. are employees or paid contractors of Integrated DNA Technologies (IDT), which sells reagents used or similar to those used in this manuscript. M.M, K.F., and R.N. are both employees of Illumina Inc. which provides a productized cloud-computing platform for doing NGS analysis. All other authors declare no conflicts of interest.

## Author Contributions

G.K. and M.M. performed all back-end bioinformatics pipeline creation. H.L. created the psnw alignment algorithm. M.M. designed all Cas9 and Cas12a guides. R.T and G.R. planned and performed all cell culture, nucleofection and library preparation. G.K performed optimization of different algorithm components and performed data analysis. N.R. performed gBlock template dilution experiments. L.T. and G.K developed code testing framework. R.N and K.F. created front-end web UI. G.K. wrote an initial draft.

---

**Figure S1.**
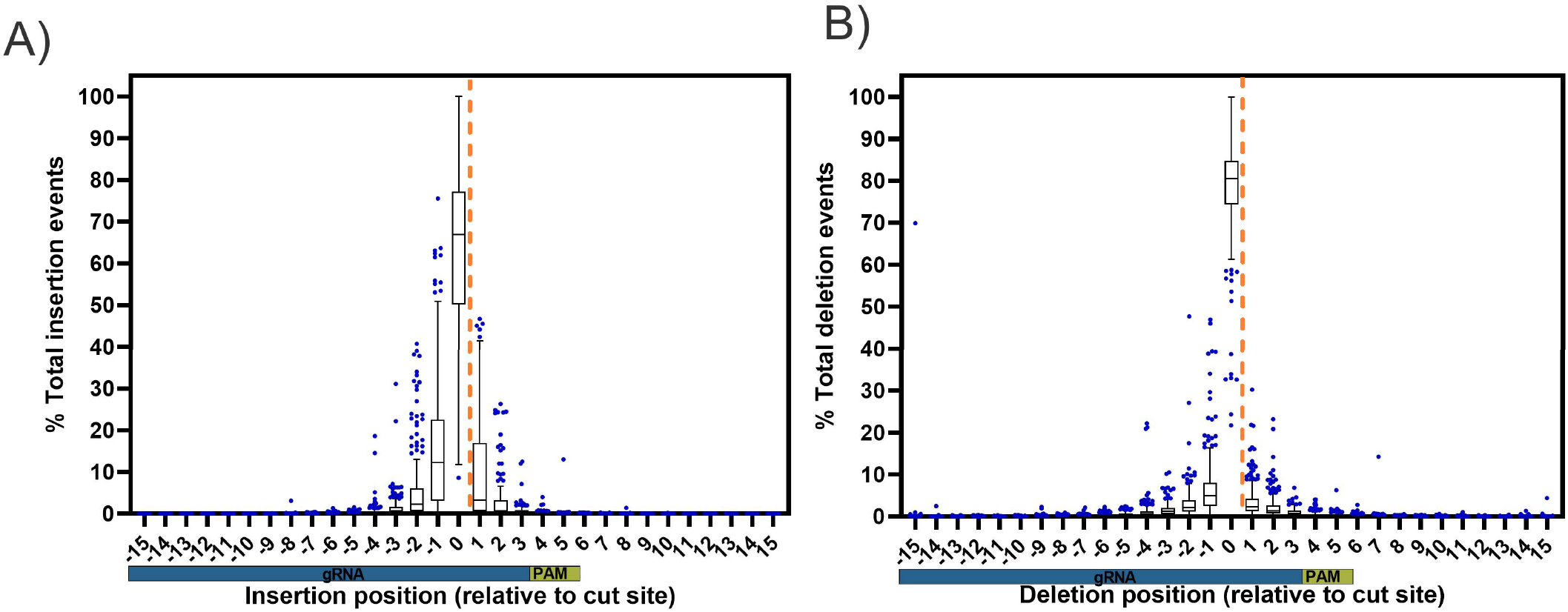
Characterization of Cas9-specific indel profiles for using the standard Needleman-Wunsch alignment algorithm (Software iteration #1). Tukey box and whisker plot of A) insertion position, and B) deletion position profiles relative to the cut site (orange dashed line) of Alt-R S.p. Cas9 V3 (n=273 guides) editing events delivered via ribonucleoprotein nucleofection into Jurkat cells analyzed using software iteration #1.

**Figure S2.**
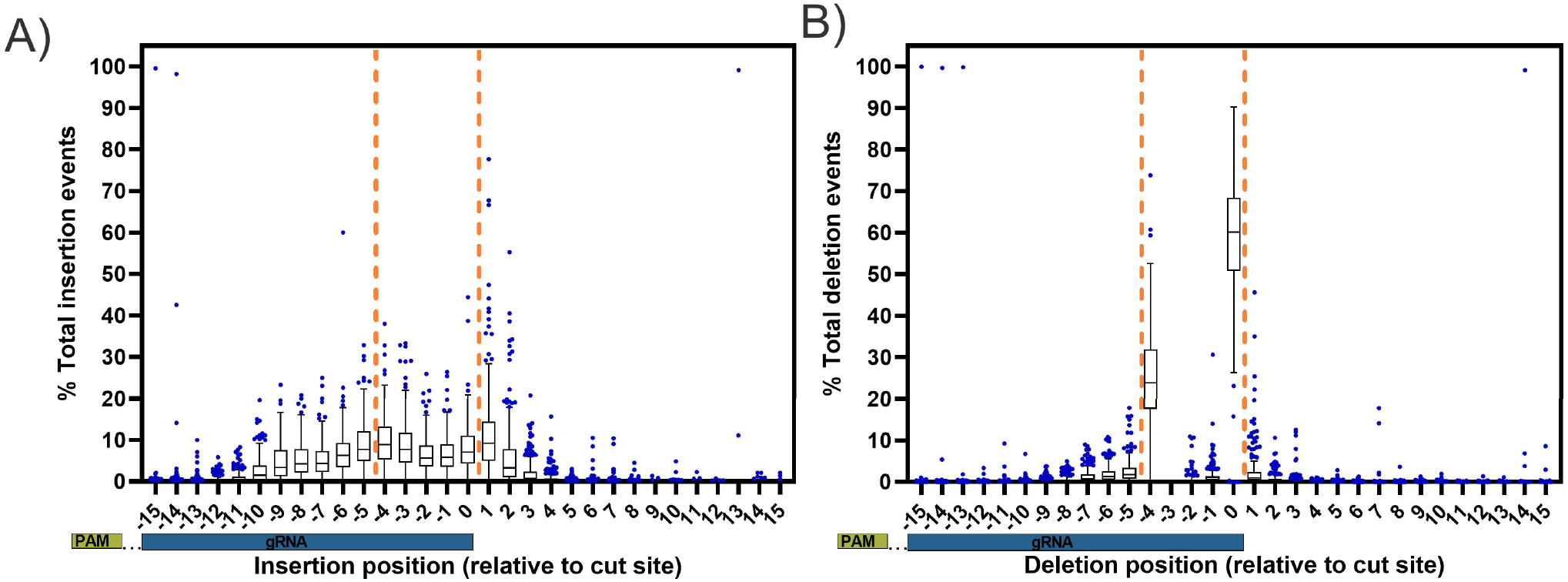
Characterization of Cas12a-specific indel profiles using the standard Needleman-Wunsch alignment algorithm (Software iteration #1). Tukey box and whisker plot of A) insertion position, and B) deletion position profiles relative to the putative nick sites (orange dashed line) of Alt-R A.s. Cas12a Ultra V3 (n=243 guides) editing events delivered via ribonucleoprotein electroporation into Jurkat cells analyzed using software iteration #1.

**Figure S3.**
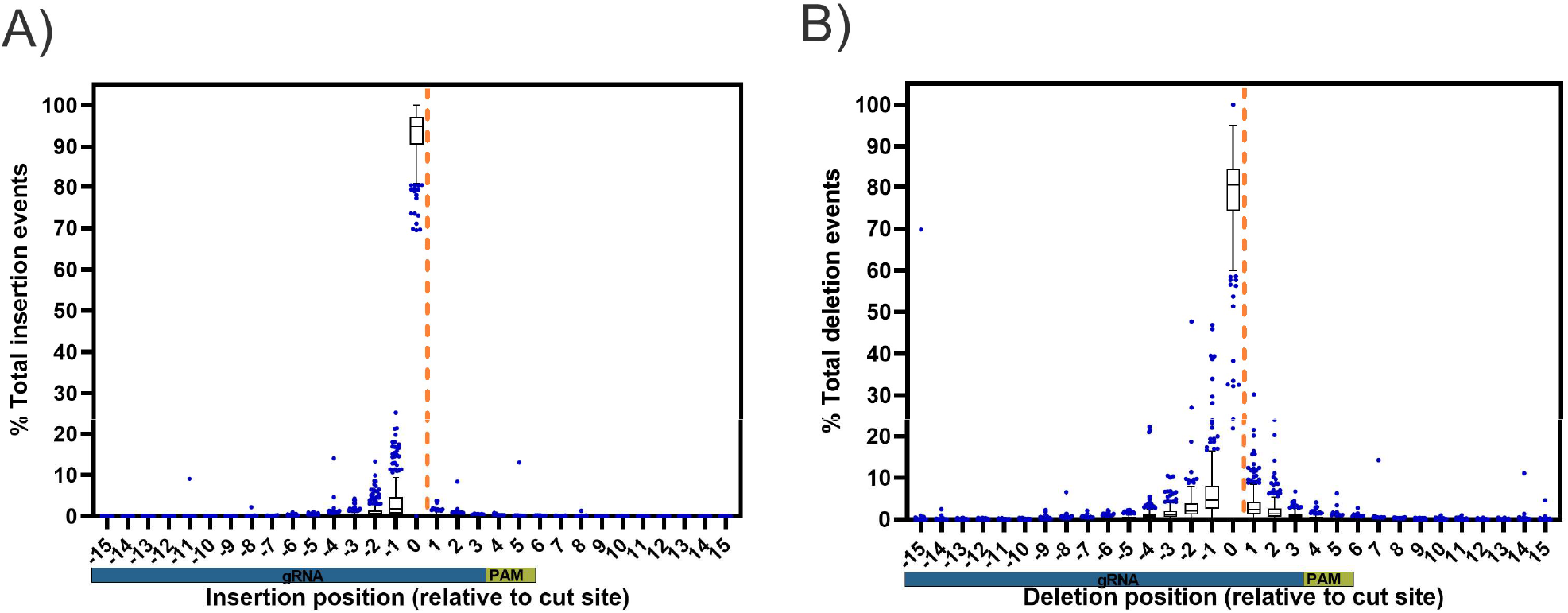
Characterization of Cas9-specific indel profiles for using psnw alignment algorithm with a single cut site bonus (Software iteration #2). Tukey box and whisker plot of A) insertion position, B) deletion position profiles relative to the cut site (orange dashed line) of Alt-R S.p. Cas9 V3 (n=273 guides) editing events delivered via ribonucleoprotein electroporation into Jurkat cells analyzed using software iteration #2.

**Figure S4.**
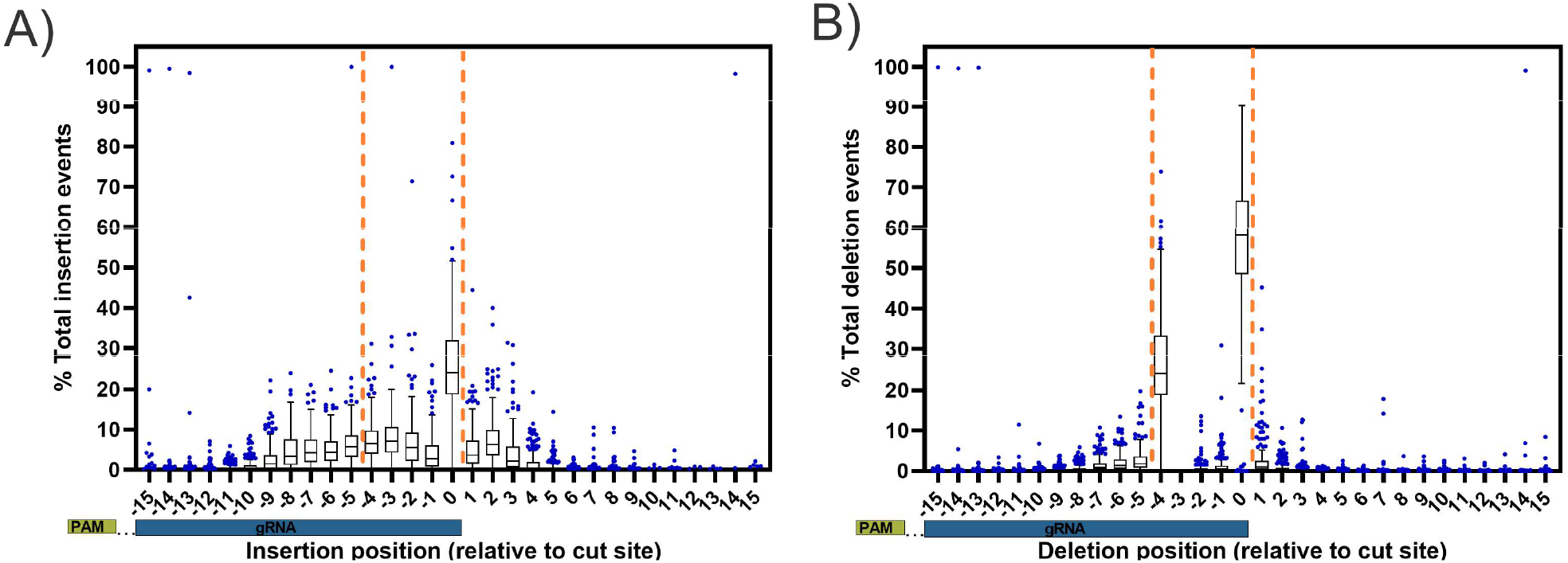
Characterization of Cas12a-specific indel profiles using psnw with a single PAM distal cut site bonus (Software iteration #2). Tukey box and whisker plot of A) insertion position, and B) deletion position profiles relative to the putative nick sites (orange dashed line) of Alt-R A.s. Cas12a Ultra V3 (n=243 guides) editing events delivered via ribonucleoprotein electroporation into Jurkat cells analyzed using software iteration #2.

**Figure S5.**
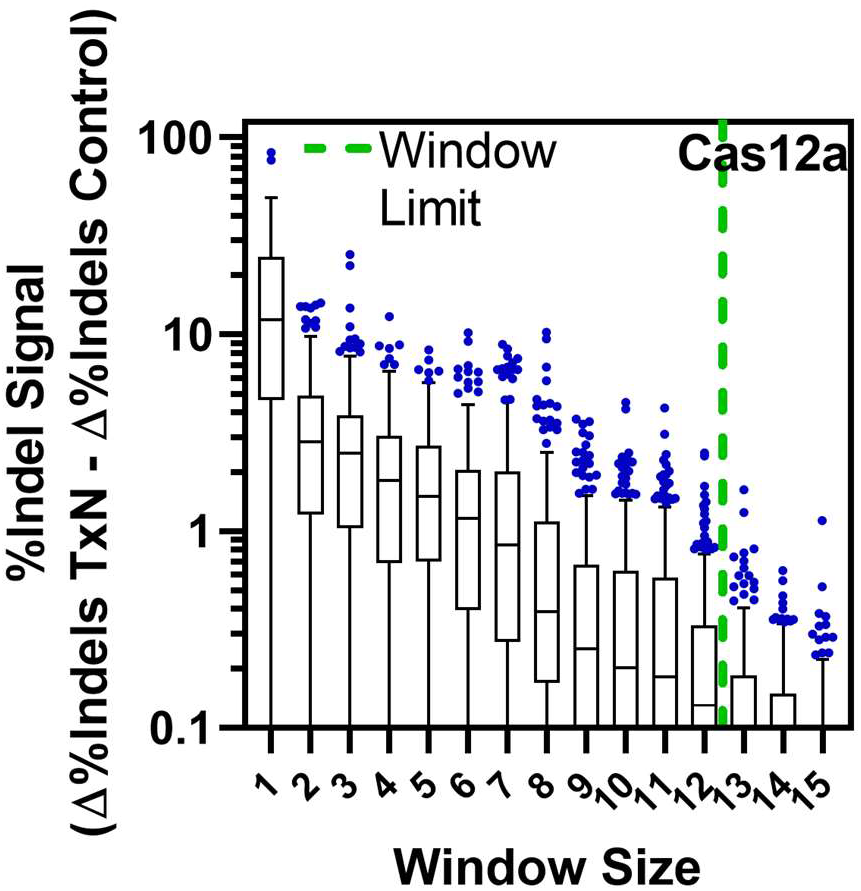
Window optimization AsCas12a centered at the PAM-distal nick site. An optimal window size (green dashed line) for annotating variants was selected for Alt-R A.s. Cas12a Ultra V3 editing in Jurkat cells at which median indel signal differences between treatment and control samples < 0.1%, with the window centered at the PAM-distal nick site.

**Figure S6.**
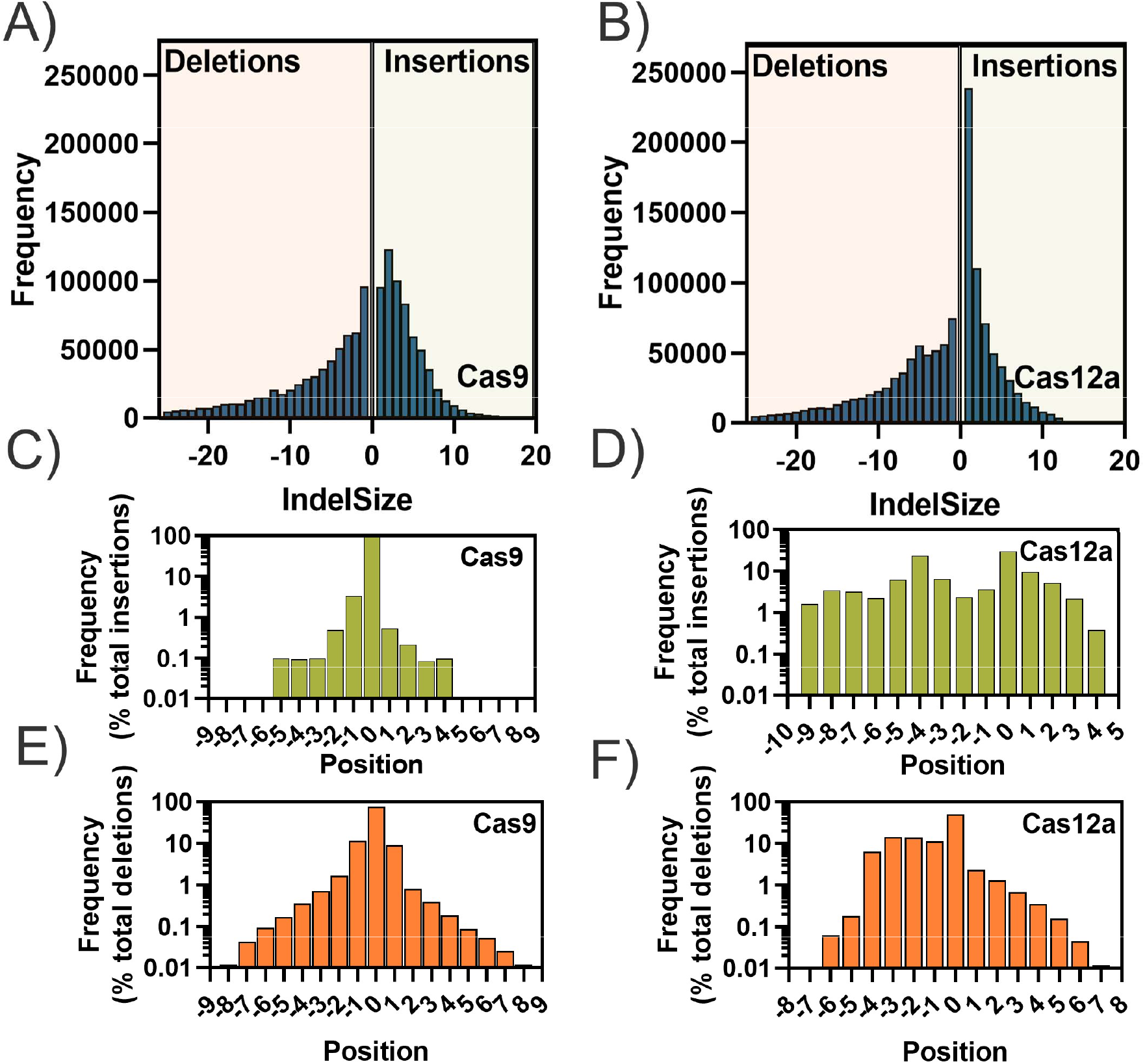
Synthetic on/off-target dataset used for pipeline validation. Characterization of synthetic CRISPR NGS on/off-target benchmarking data A-B) indel sizes, C-D) insertion positions, and E-F) deletion positions, all modeled based on experimental Alt-R S.p. Cas9 V3 or Alt-R A.s. Cas12a Ultra V3 editing data in Jurkat cells.

**Figure S7.**
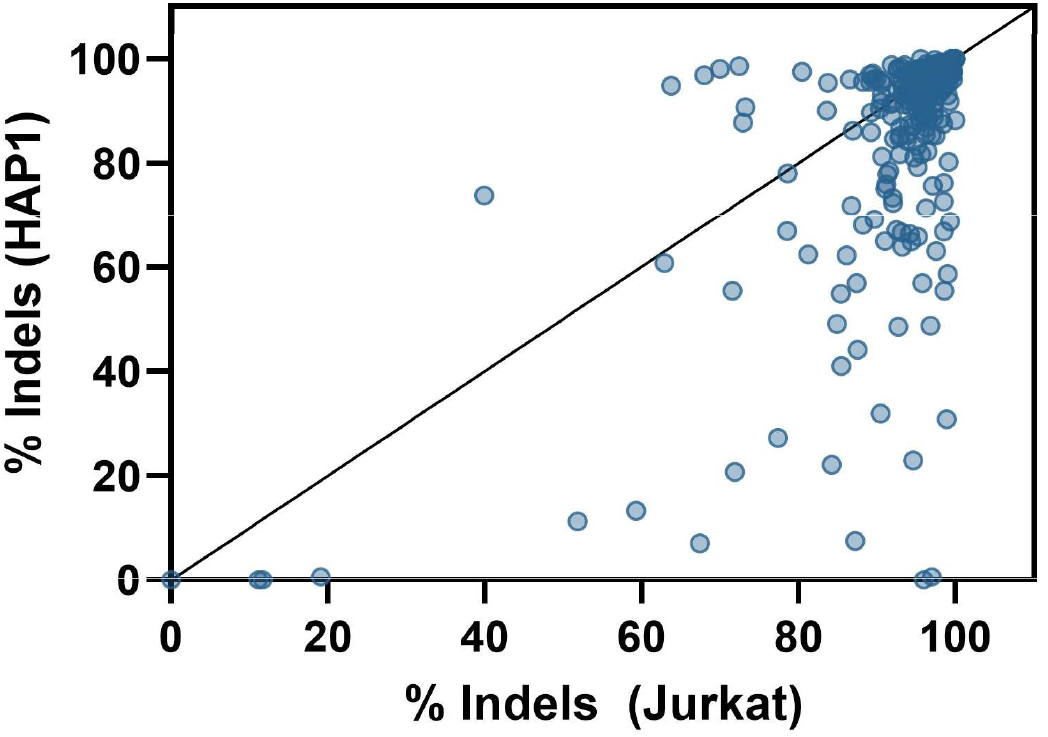
HiFi Cas9 editing in HAP1 and Jurkat Cells. Quantification and comparison of indel editing by CRISPAltRations in HAP1 and Jurkat cell lines with Alt-R S.p. Cas9 V3 delivered via ribonucleoprotein (n=273 unique gRNAs).

**Figure S8.**
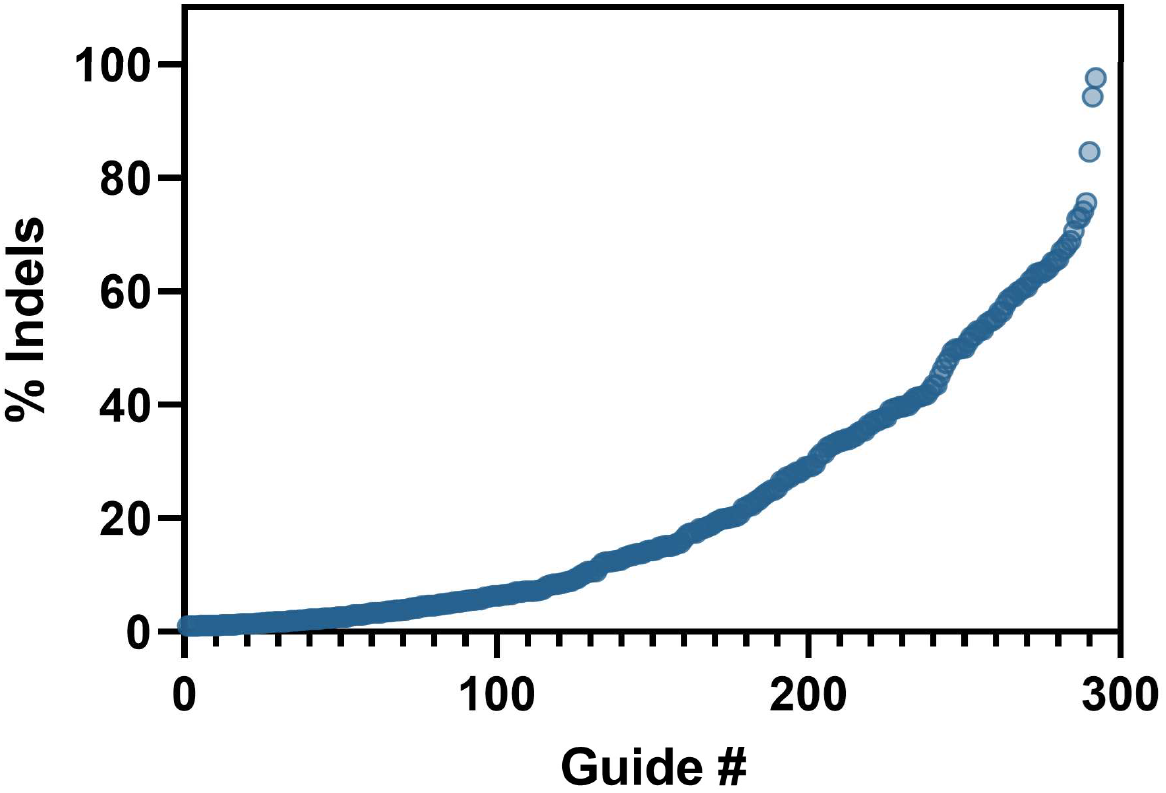
AsCas12a editing efficiency in Jurkat. Quantification of editing by CRISPAltRations in Jurkat delivered Alt-R A.s. Cas12a Ultra V3 via ribonucleoprotein electroporation at a suboptimal concentration (n=243 unique gRNAs)

**Figure S9.**
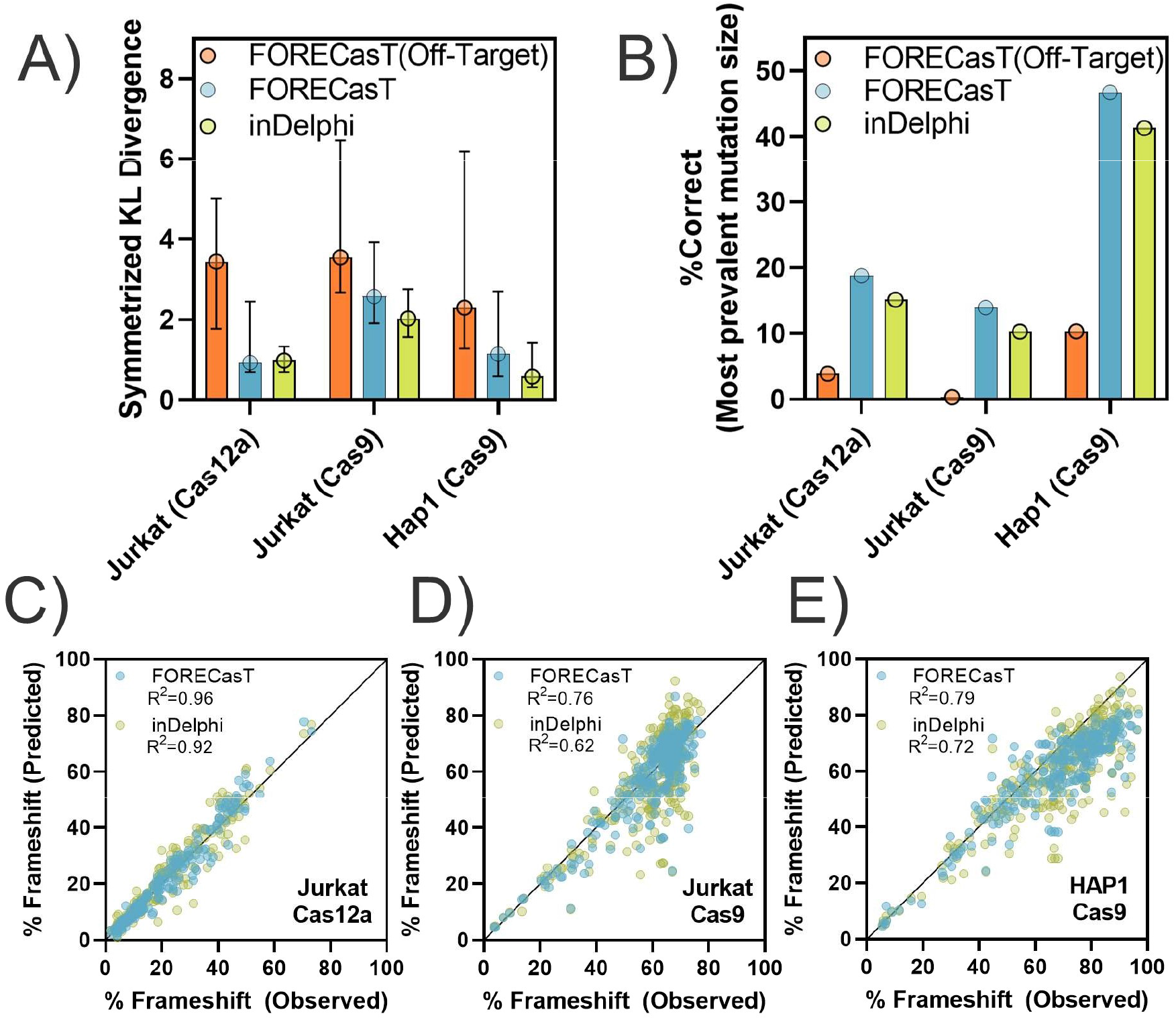
Performance of *in-silico* mutation profile prediction tools. FORECast and inDelphi were evaluated for the ability to predict mutation size distributions similar to what was observed Jurkat/HAP1 cells treated with Alt-R S.p. Cas9 V3 Cas9 or Alt-R A.s. Cas12a Ultra V3 by measuring A) symmetrized KL divergence between observed and predicted profiles (median +/- IQR) and B) the mean accuracy predicting the most prevalent mutation. Linear regression was performed using predicted vs observed frameshift frequencies for C) Jurkat + Cas12a D) Jurkat + Cas9 and E) HAP1 + Cas9 treated cells with a line of identity (solid black line) displayed at y = x.

**Figure S10.**
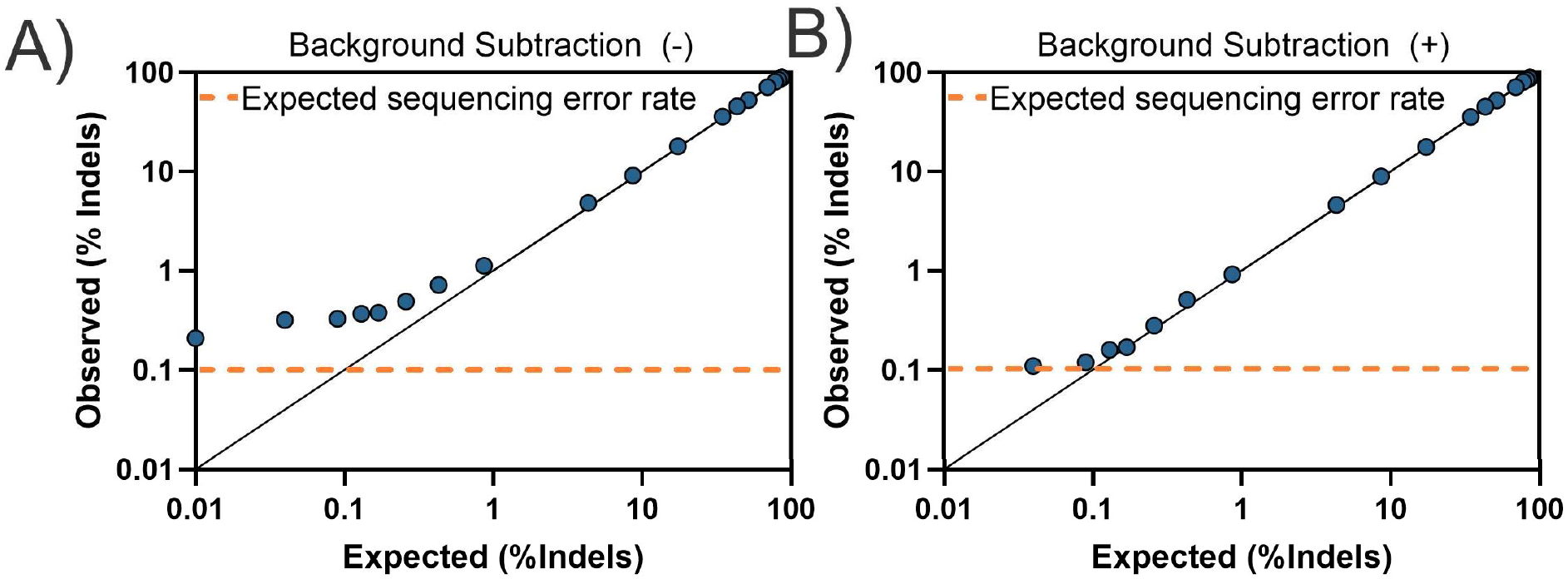
Pipeline indel detection sensitivity. Pipeline indel detection concordance (black line) with a titrated mixture of gBlocks with known concentrations of indels for an HPRT target (>40,000 reads per sample) sequenced with MiSeq v3 chemistry A) before and B) after a simple background subtraction, performed by subtracting the percent indels observed in an unmodified gBlock control from all samples.

**Figure S11.**
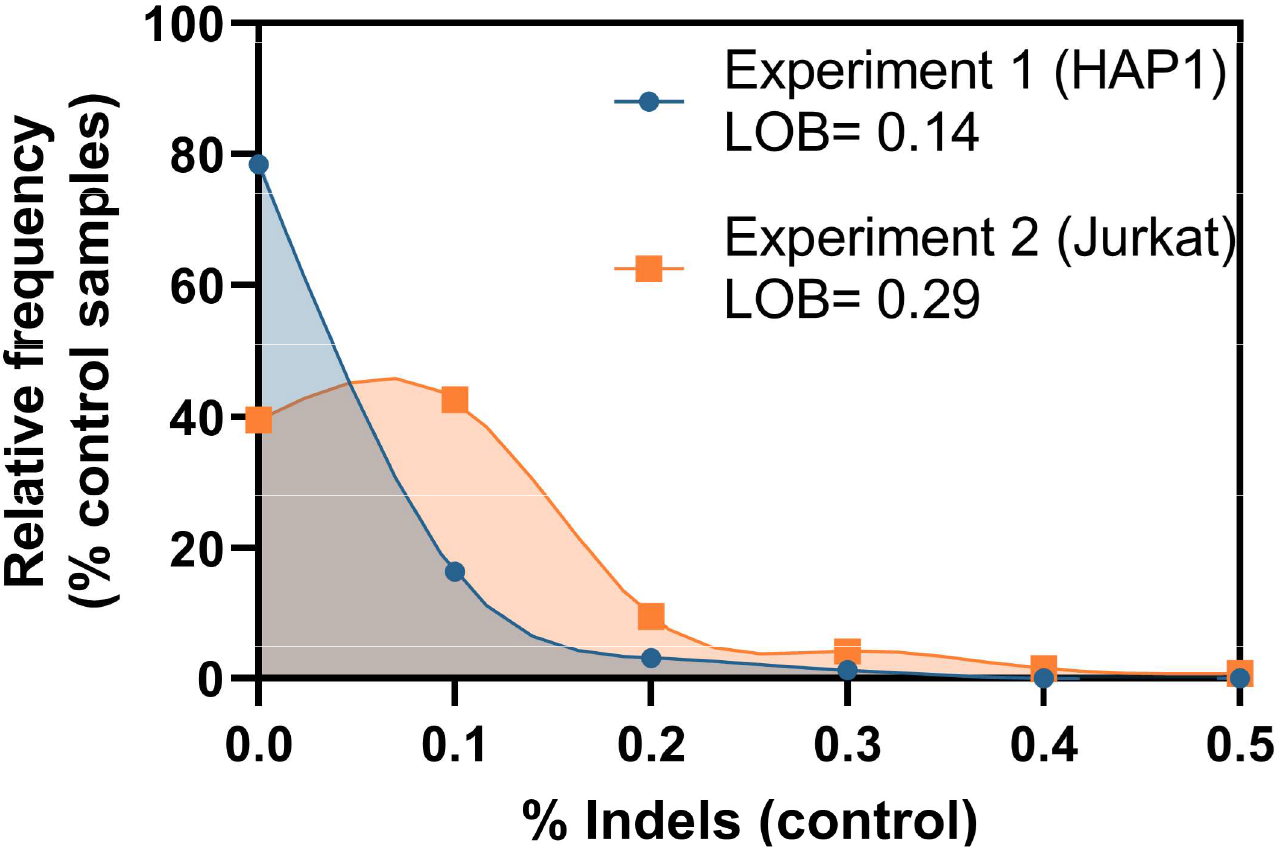
Evaluation of indel background noise in two experiments. Relative frequency of unedited control samples with variable indel editing signal (binned in 0.1% intervals) for the same genomic targets from Jurkat (n=260) and HAP1 (n=158) cell lines in two separate experiments with high read depth (> 10,000 read pairs). Limit of Blank; LOB

**Figure S12.**
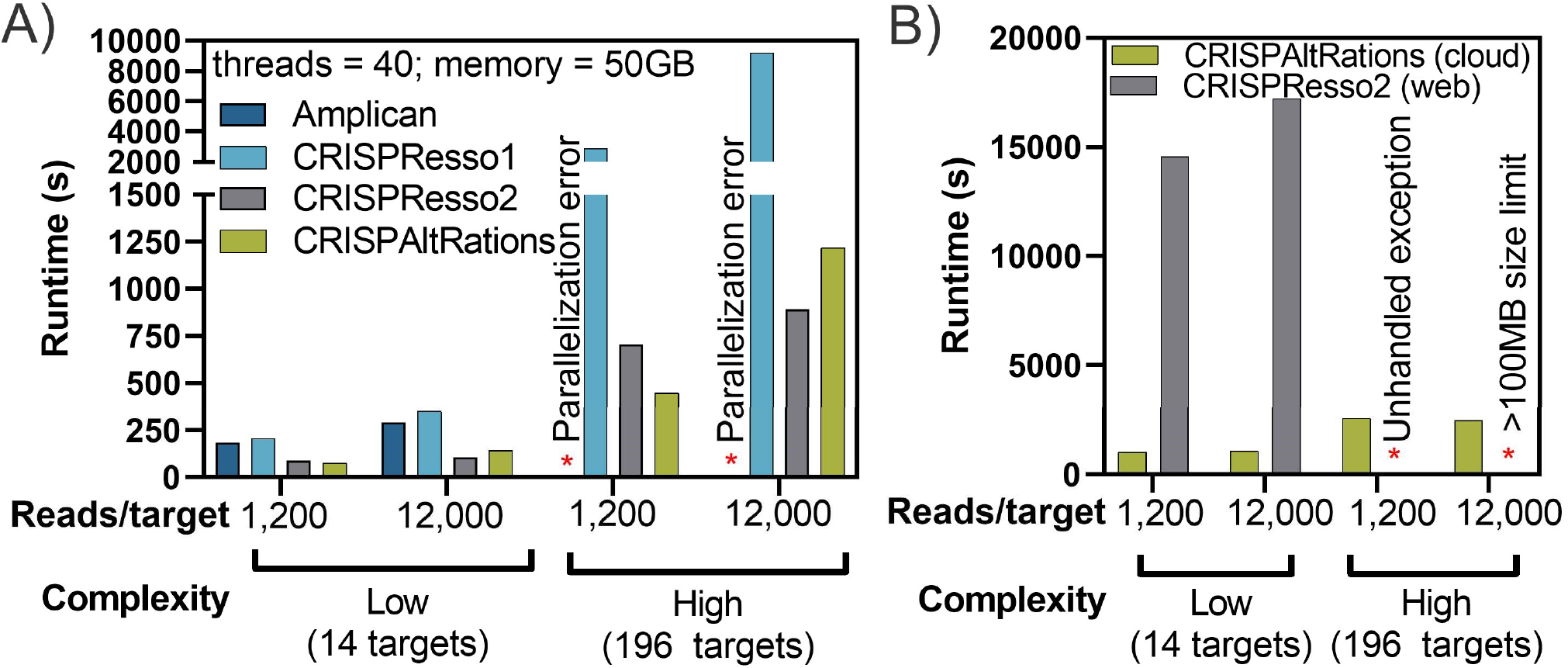
Pipeline runtime requirements. All multiplex compatible pipelines were ran against synthetic multiplex on/off-target datasets with 14 or 196 targets at varying read depth. Runtime in seconds was recorded for A) Command line interface and B) Web UI runs. Runs that failed submission or analysis are indicated (*).

